# Osteopontin promotes lesion repair during *Staphylococcus aureus* skin infections

**DOI:** 10.64898/2026.03.19.712901

**Authors:** Emily E Neville, Mari L Shinohara, Jennifer Y Zhang, Abhay PS Rathore, Soman N Abraham

## Abstract

Skin infections are among the most common types of infections in the United States and pose a serious health threat. Effective resolution requires both bacterial clearance and tissue repair, however the mechanisms underlying tissue repair remain poorly understood. Here, we investigate the cellular responses in a murine model of *S. aureus* dermonecrotic skin infection using single cell transcriptomics. Within the infected lesions, we identified a major influx of heterogeneous neutrophils, including a distinct subtype expressing high levels of *Spp1*, the gene encoding osteopontin (OPN). Cell-cell interaction analysis identified *Spp1* signaling as the most enriched pathway in infected tissue. In OPN knockout (KO) mice, transcriptomic profiling revealed a lower proportion of fibroblast and keratinocyte cells at the infection site, and a reduction in associated repair signaling pathways. *In vivo*, OPN KO mice developed significantly larger lesions during skin infection, whereas WT mice treated with recombinant OPN showed accelerated healing relative to untreated controls. Together, these findings highlight the importance of specific neutrophil subtypes and OPN signaling as critical mediators in infected tissue repair. Further our study suggests that modulating OPN may represent a promising therapeutic strategy to accelerate infected tissue repair.

## Introduction

Skin and soft tissue infections (SSTIs) are an abundant problem in the US impacting 14 million people annually [1]. These infections pose a significant health threat as they can progress into deeper tissue infections such as cellulitis, abscess, and even sepsis [2, 3]. *Staphylococcus aureus* (*S. aureus*) is the most common cause of SSTIs in the US [4]. These infections can be particularly challenging to treat as *S. aureus* α-hemolysin toxin causes dermonecrosis [5] - increasing tissue damage which can lead to delays in wound closure and subsequent infection recurrence [6].

During SSTIs, bacteria enter the skin through a breach in the skin barrier where they can multiply and cause infection [7]. The host then undertakes both immune responses to combat the bacteria as well as tissue repair processes to close the lesion [8]. Early during *S. aureus* SSTI, there is robust recruitment of neutrophils to the infection site [9]. These neutrophils are critical for infection control as they phagocytose and kill bacteria via oxidative bursts. They also prevent the spread of infection by releasing NETs and creating abscesses to wall off the bacteria that cannot be immediately killed [8]. This is followed up by later infection responses including the recruitment of IL-17 producing γδ T cells which help to mediate bacterial clearance [10, 11]. In the skin during sterile wound repair, neutrophils are cleared early on and repair is mainly mediated by other cell types [12]. Macrophages and fibroblasts are critical for matrix deposition and ECM remodeling while keratinocytes expand and migrate to promote reepithelialization to repair the skin breach [13]. While there is a breadth of knowledge on how tissue repair occurs during sterile wounding, less is known about how repair occurs and what cell types and molecules are critical to repair during infection.

Single-cell RNA sequencing (scRNA-seq) is a powerful technology to identify cell types, cellular heterogeneity and predict functions of different cells based on changes in gene expression [14]. Through this technology, researchers have begun to identify subsets of neutrophils that have different characteristics and non-classical functions [15]. For example, a population of pro-angiogenic neutrophils have been found in both inflammation and cancer [16, 17]. Subsets with antigen presenting phenotypes have also been shown to be highly enriched during cancer [18, 19]. A neutrophil subset expressing high levels of interferon stimulated genes has been found to expand during infections [20]. Additionally, neutrophils roles and functions can change based on what tissue they are found in, gaining different phenotypes as they move from bone marrow (BM) or blood to various tissue types [21]. Neutrophils have previously been found to have a repair function during muscle injury [22]. They have also been found to cleave collagen and promote remodeling to allow for cell migration critical to repair [23]. Based on the wide variety of neutrophil responses observed across tissue types and disease conditions, it is conceivable that neutrophils during SSTI could also have a role in the repair process.

In this study, we used a dermonecrotic *S. aureus* skin infection model combined with single cell transcriptomics and genetic knockout approaches to identify key regulators of infected tissue repair. We discovered that neutrophils overwhelmed the infection site, and that neutrophils recruited to the skin were reprogrammed to have heterogeneous functions outside of classical bacterial killing. We found a subset of neutrophils expressed high levels of *Spp1,* a mediator which was the top upregulated pathway during infection. Osteopontin (OPN), encoded by *Spp1,* is a glycophosphoprotein which can be secreted during inflammation and infection. We found that OPN plays a critical role in promoting tissue repair following skin infection. Notably, treatment with recombinant OPN accelerated infected lesion healing, highlighting its potential as an effective therapeutic strategy to enhance recovery from SSTIs.

## Results

### Temporal dynamics of *S. aureus* abscess formation and immune response during skin infection

To address the gap in knowledge on the progression of *S. aureus* SSTIs and how different cell types contribute to the temporal dynamics of bacterial skin infections, we used a dermonecrotic *S. aureus* infection model on which we have previously published [24]. We chose this model because it recapitulates many of the features seen clinically during *S. aureus* skin infections including rounds of scabbing, biofilms, abscess formation, and eventual reepithelialization [25, 26], enabling the study of both bacterial clearance and tissue repair phases during infection.

Mice were intradermally infected with *S. aureus* complexed with cytodex beads. Skin samples at the infected lesion site were collected at 1-, 7- and 14-days post infection (DPI) for assessment (Fig. 1a). We chose these timepoints to represent the various stages of infection from a closed inflamed lesion with early immune recruitment at 1 DPI, to the peak of immune response and start of repair at 7 DPI followed by reepithelialization of the lesion at 14 DPI (Supplemental Fig 1a). To assess how the bacterial load changed in our model over time we performed CFU assays. Interestingly, bacterial load remained stable for 7 days before, dropping off at 14 DPI and resolving at 19 DPI (Fig. 1b). This persistence may be due to the abscess walling off the infection, preventing bacterial clearance. Consistent with this, scanning electron microscopy (SEM) at 7 DPI revealed bacteria embedded within a biofilm matrix at the abscess site (Fig. 1c), a hallmark of complicated *S. aureus* SSTIs [27, 28].

**Figure 1.**
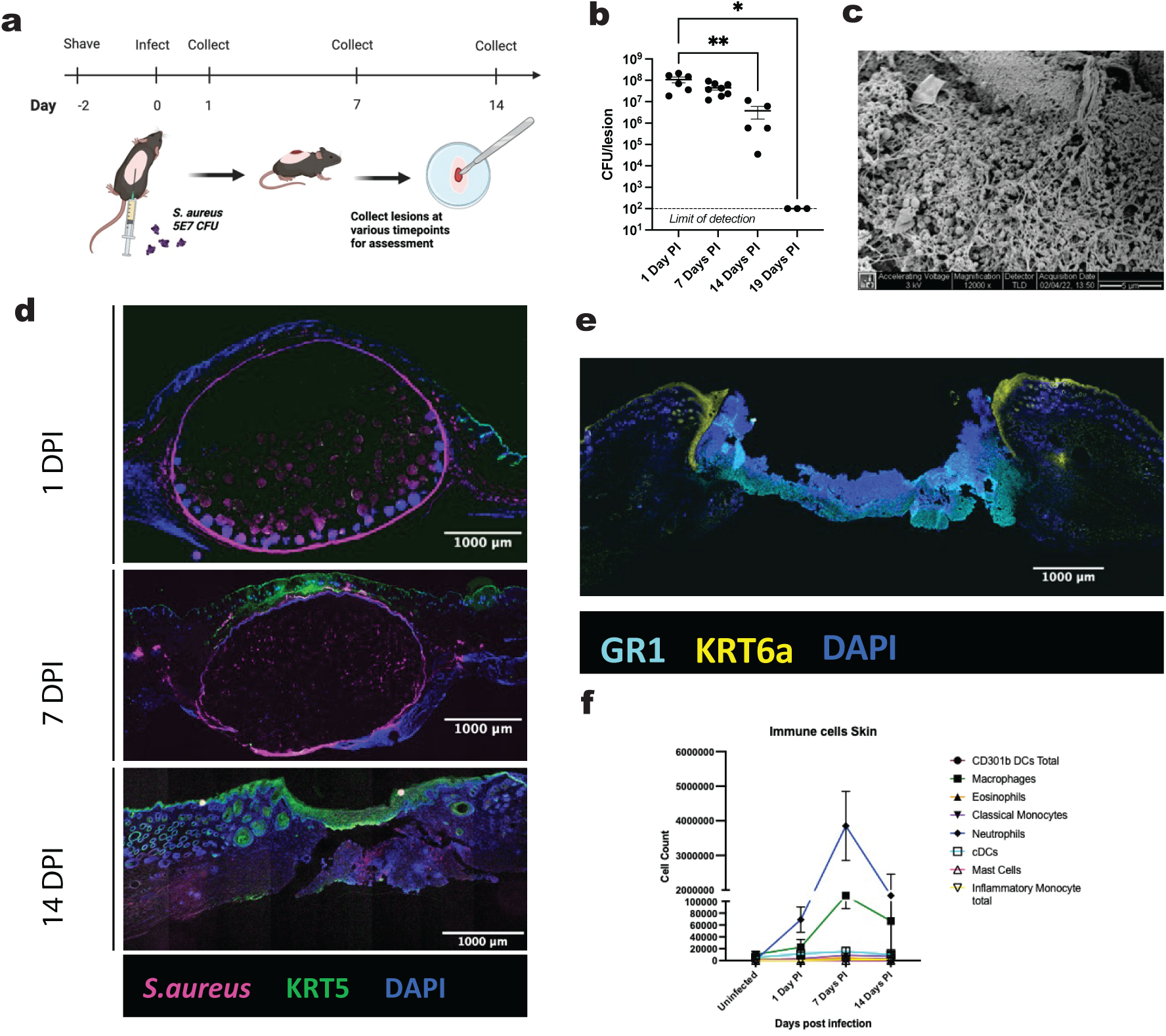
*S. aureus* skin infection response is dominated by neutrophil influx. (a) Timeline of skin infection protocol, WT mice had hair removed 2 days prior to intradermal injection of 5E7 *S. aureus* complexed with cytodex beads. Skin was then collected at various timepoints to assess the infected lesion (b) Bacterial load in the skin was assessed at 1-, 7-, 14- and 19-days post infection via CFUs (n = 3-8 mice per group, from 2 or more independent experiments) (c). SEM imaging of infected lesion site at 7 DPI, representative of 3 mice from 2 independent experiments, scale bar = 5 μm. (d) Representative tilescan images of immunostaining at lesion site at 1, 7 and 14 DPI. Sections were stained for *S. aureus* (magenta), KRT5 (green) and DAPI nuclear stain (blue), scale bar = 1000 μm (n = 3-4). (e) Representative tilescan images of immunostaining for Neutrophil Gr-1 (cyan), KRT6a (yellow) and DAPI nuclear stain (blue) at 7 DPI scale bar = 1000 μm (n = 3). (f) Flow cytometry cell counts of innate immune cell populations in skin from uninfected and infected mice at indicated timepoints post infection, n = 4-5 mice. Statistical analyses were performed with one-way analysis of variance (ANOVA) where there were > 2 groups with Tukey’s post hoc test. Means ± SEM, \**P* < 0.05, \*\**P* < 0.01. PI, post infection.

To visualize the dynamics of infection and lesion repair, we employed immunofluorescence microscopy. At 1 DPI, we observed the formation of an abscess-like structure of bacteria and beads within the skin (Fig. 1d). The abscess persisted through the 7-day timepoint and resolved by 14 DPI, though IF staining showed that bacteria had not fully cleared in the lesion consistent with our CFU findings. IF staining also showed the loss of KRT5, a marker of basal epithelial cells, at 1 DPI above the abscess site, likely due to dermonecrosis of the epithelial cells above the infection. At 7 days post infection we see KRT6a expression in the edge epithelium growing around the lesion site (Fig. 1e) this is a marker seen in migrating wound edge cells [29] indicating that tissue repair has begun to occur at this timepoint. We can see significant repair though with reepithelialization at the site indicated by the KRT5+ basal keratinocytes spanning the lesion site by 14 DPI (Fig. 1d). This data shows that repair can occur despite the ongoing presence of bacteria in the lesion.

To further our understanding of the dynamics of immune responses during *S. aureus* infection, we used flow cytometry to assess immune cell recruitment over time. Concurrent with what is seen in human skin infections, neutrophils dominated the infection response [7] making up roughly 90% percent of CD45+ cells at the lesion site by 7 DPI (Fig. 1f & Supplemental Fig. 1c, 1d). This corresponded with the peak level of *Il1b* (Supplemental Fig. 1b), a known inflammatory marker that has previously shown to be elevated during skin infection [30]. To confirm the presence of neutrophils at the infection site, we did immunofluorescence microscopy which show accumulation of Gr1+ neutrophils under the open lesion site at 7 DPI (Fig. 1e). We also observed increasing levels of macrophages at 7 DPI (Fig. 1f) which are known to promote bacterial clearance as well as tissue repair [31]. From the adaptive immune compartment, we observed an increase in IL-17+ CD4 T cells at 14 DPI (Supplemental Fig. 1e). IL-17 is known to be critical in resolution of *S. aureus* skin infections and can promote the continued recruitment of neutrophils into the infection site [10]. Together these data define skin infection responses over time revealing the predominance of the neutrophil response, and the temporal dynamics as SSTI response transitions from bacterial clearance to tissue repair.

### Transcriptomic analysis of cells from infected tissue reveals heterogeneous neutrophil subsets with distinct functional roles

To further investigate the roles of cells responding to *S. aureus* skin infection, we took an unbiased scRNA-seq transcriptomic approach. We selected the 7 DPI timepoint to investigate, as it was when we saw high bacterial numbers, a distinct abscess, and peak immune cell recruitment, as well as signs of tissue repair as described in Fig. 1. Lesion site samples from uninfected mice and mice at 7 DPI were collected, dissociated for single cells and subjected to single cell transcriptomic analysis using the 10X Genomics platform (Fig. 2a). Following filtering for high quality cells, we identified several immune and nonimmune epithelial cell types across our infected and healthy tissue (Fig. 2b, 2c) based on differentially expressed marker genes (Fig. 2d).

**Figure 2.**
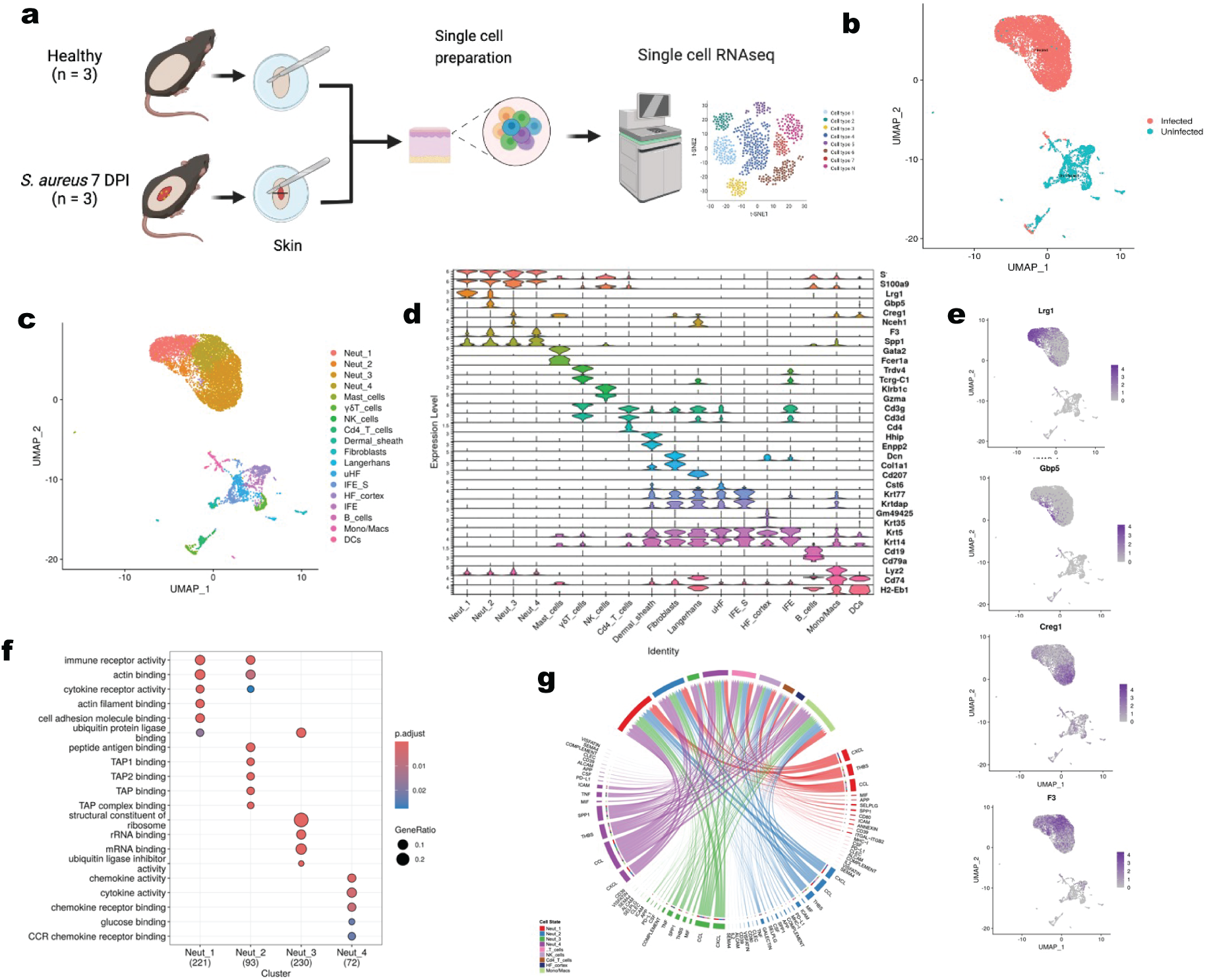
scRNA-seq of *S. aureus* infected skin reveals diverse neutrophil populations. (a) Schematic for scRNA-seq, skin from each group of mice (uninfected) and mice 7 days post *S. aureus* infection was dissociated, sorted for live cells and sequenced using 10x Genomics (n = 3 mice per group, pooled) (b) UMAP dimensional reduction of infected and uninfected samples. (c) UMAP of scRNA-seq skin cells showing 18 cell populations (d) Violin plot of gene expression levels for representative genes used to identify cell type for each cell population identified in (c). (e) Marker gene expression of neutrophil subpopulations on UMAP (f) Gene ontology (GO) enrichment analysis of upregulated activity for neutrophil subpopulations (g) CellChat chord diagram showing all signaling pathways originating from neutrophil subpopulations in infected mice.

To investigate how skin tissue composition changes during infection, we first characterized the cellular landscape in healthy skin tissue. Epithelial cells were the dominant cell type in skin from uninfected mice. This was expected as there are limited numbers of immune cells present in the skin at steady state. Epithelial cell types were defined using previously described marker genes [32–34]. We identified several hair follicle cell types including upper hair follicle keratinocytes (uHF) expressing *Cst6*, HF cortex cells expressing *Krt35* and *Gm49425* (*Krt85*), intrafollicular epithelial cells (IFE) expressing *Krt5* and *Krt14*, and suprabasal IFEs (IFE_S) which additionally expressed high levels of *Krt77* and *Krtdap* (Fig. 2d).

In contrast to healthy skin, the infected skin samples were dominated by immune cell populations, with neutrophils comprising the largest group in our single cell dataset (Fig. 2c). Recent studies have shown that neutrophils are not a homogeneous population, but instead exhibit diverse phenotypes and functions [15]. Given their abundance during *S. aureus* infection, we investigated whether these neutrophils represented distinct subsets and what functional roles they might play. Specifically, we were interested in identifying neutrophil subsets involved in processes beyond classical bacterial clearance, such as tissue repair, particularly since both bacterial elimination and tissue regeneration are actively occurring at 7 DPI. Indeed, looking at our data, we identified four distinct clusters of neutrophils expressing *S100a8* and *S100a9* markers (Fig. 2d), revealing substantial heterogeneity in the neutrophil compartment during infection. The Neut1 cluster expressed high *Lrg1* (Fig 2e) and was characterized by high immune receptor, cytokine and actin binding (Fig. 2f). Additionally, CellChat signaling analysis showed increased ANNEXIN & CD80 signaling (Fig. 2g), both of which have been shown to promote inflammation resolution and tissue repair. Together these indicate that the Neut1 cluster likely plays a role in tissue repair during infection.

Neut2 cluster, characterized by high levels of *Gbp5* (Fig 2e), was shown to have significant signaling of the transporter associated with antigen processing (TAP), TAP1 and TAP2 binding pathways (Fig. 2f). The Neut2 cluster also contributed highly to CD80 signaling (Fig 2g, Supplemental Fig. 2c) which is traditionally highly expressed in APCs and can promote T cell signaling [35, 36]. Together, these data indicates that the Neut2 cluster may have a key role in antigen presentation of *S. aureus* to adaptive immune cells. Antigen presentation has been shown to be critical for the development of IL-17+ γδ T cells in the draining lymph nodes leading to eventual bacterial clearance [11]. We observed high *Il17a* and *Il17f* expression in γδ T cells from infected mouse skin (Supplemental Fig. 2b). Additionally, the Neut2 cluster had a predominant sender role in Galectin signaling (Fig. 2g, Supplemental Fig. 2d). This was largely mediated through *Lgals9* signaling (Supplemental Fig. 2e). *Lgals9* has previously been shown to promote keratinocyte proliferation in IL-17+ environments [37], indicating that Neut2 cells could also have a pro tissue repair role.

The Neut3 cluster, marked by *Creg* expression (Fig. 2e), had increased pathways associated with a classical role in bacterial killing. These included enriched TNF, Complement, and CXCL pathways (Fig. 2g), as well as increased KEGG signaling related to the phagosome, lysosome, and oxidative phosphorylation (Supplemental Fig. 2a). The Neut4 cluster exhibited significant GO enrichment for cytokine and chemokine activity (Fig. 2f, 2g), suggesting a specialized role in immune signaling. This subset showed the highest levels of CCL signaling among neutrophil populations (Fig. 2g), indicating active participation in chemokine-mediated communication. KEGG pathway analysis further revealed enrichment in Toll-like receptor signaling, C-type lectin receptor signaling, and NF-κB signaling pathways (Supplemental Fig. 2a), all of which are critical for initiating and regulating inflammatory responses [38–40]. Collectively, these data suggest that Neut4 neutrophils are key contributors to cell signaling and immune modulation during *S. aureus* skin infection.

Other immune cells present in our infected dataset including a monocyte/macrophage population, dendritic cells (DCs), natural killer (NK) cells, CD4+ T cells, and *IL17* expressing γδ T cells (Fig. 2c, 2d, and Supplemental Fig. 2b). IL-17 is known to be highly expressed during skin infections and is a key cytokine in promoting skin infection responses [41]. The high levels of *Il17a* and *Il17f* observed could help to promote the drop off in bacterial levels that we observed from the 7- to 14-day timepoint in Figure 1b.

Together, these analyses provide insight into the transcriptomic landscape at a critical timepoint during *S. aureus* skin infection, when both bacterial clearance and tissue repair are actively occurring. Our data reveal a robust neutrophil response within the infected tissue. Strikingly, these neutrophils were heterogenous, comprising distinct subsets with predicted functional roles in antigen presentation, cell signaling, and tissue repair which together highlight the multifaceted contributions of neutrophils beyond their classical bacteria-clearing functions.

### Enhanced SPP1 signaling by recruited neutrophil subsets during skin infection

To further investigate the roles of immune cells recruited to the lesion site, we performed receptor ligand analysis to uncover the signaling pathways between cells in our scRNA-seq data. Using CellChat analysis [42], we identified 17 signaling pathways that were significantly enriched during *S. aureus* infection at 7 DPI (Fig. 3a). While we saw increases in expected pathways that have key roles during infection like CCL, CXCL, and complement, we were surprised to find that the topmost enriched pathway was SPP1. *Spp1*, which encodes the glycophosphoprotein osteopontin (OPN), is implicated in a wide range of physiological and pathological processes across various disease contexts [43]. OPN can exist in a secreted or intracellular form [44, 45]. Secreted OPN has been shown to be a potent chemoattractant. Additionally, it can inhibit apoptosis thereby contributing to cell survival in different disease contexts [46]. OPN is expressed by a wide range of cell types, including leukocytes and fibroblasts, and has been intensively studied in macrophages [43, 47]. While we did observe *Spp1* signaling from the mono/mac population, neutrophils were unexpectedly the highest producers of *Spp1* (Fig. 3b). Neutrophils, were previously reported to have little to no *Spp1* expression [48], however during skin infection, we discovered that a cluster of neutrophils, Neut4, were the top producers of *Spp1* with limited *Spp1* also produced by the Neut3 cluster (Fig. 3b). The Neut4 population was predicted to express OPN to interact with multiple receivers through CD44 as an OPN receptor (Fig. 3b-d). CD44 is known for its role in adhesion, activation and proliferation of various cell types [49]. IF co-staining for the neutrophil marker Gr-1 and OPN showed high levels of OPN in the neutrophilic band surrounding the lesion site (Fig. 3e). This co-localization supports the idea that OPN is concentrated in areas of the lesion heavily infiltrated by neutrophils.

**Figure 3.**
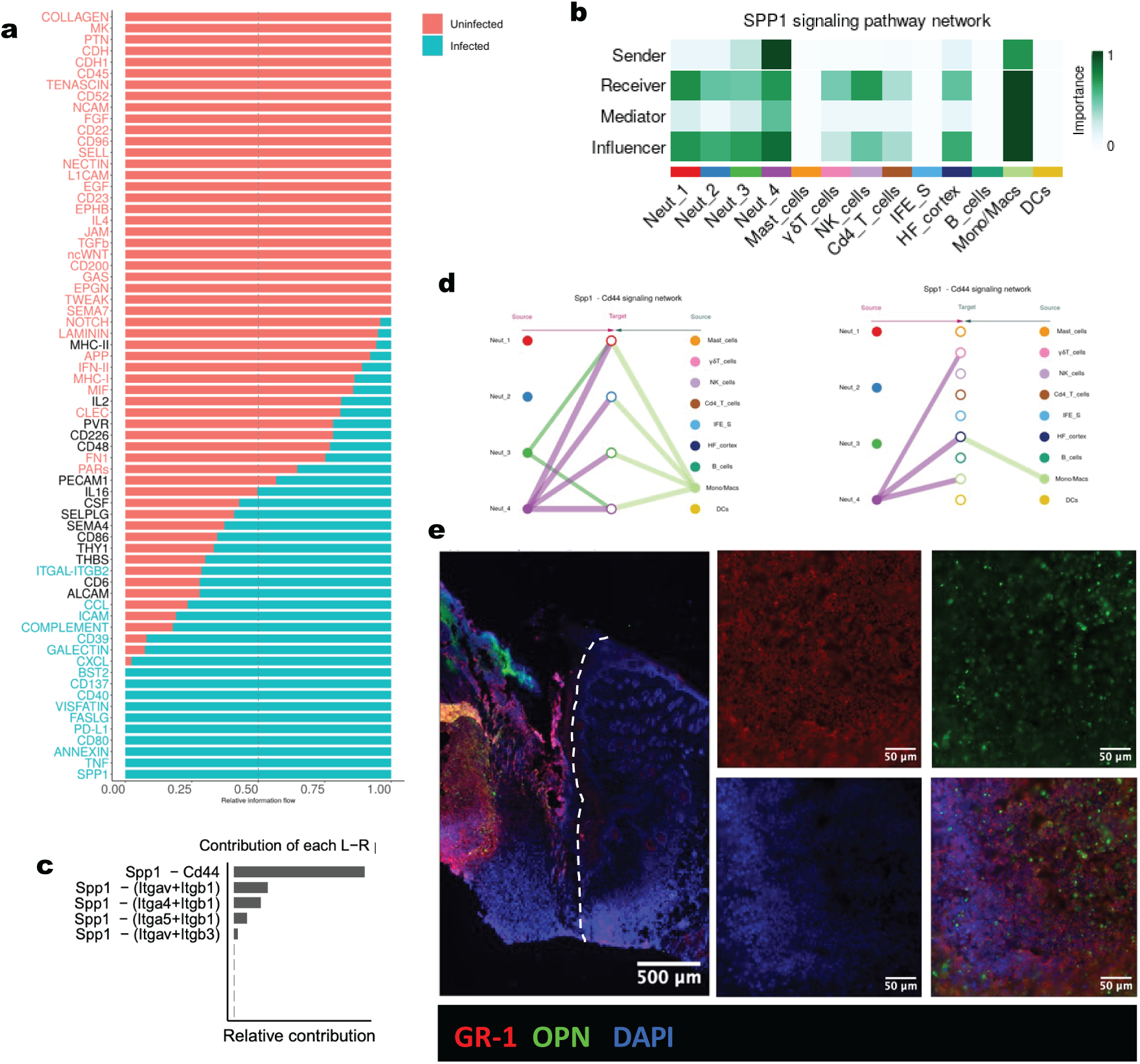
Receptor ligand analysis reveals SPP1, the top upregulated signaling pathway during infection, is strongly expressed in a subset of neutrophils. (a) CellChat analysis of intercellular communication networks in uninfected (pink) and infected (blue) WT mice. Pathways where the text is blue or pink represent pathways that are significantly upregulated in the infected or uninfected skin, respectively. (b) Heatmap of SPP1 signaling pathway network in infected skin showing the dominant cell types predicted to be involved in SPP1 signaling based on gene expression. (c) Contribution of each ligand receptor pair in the SPP1 signaling network to the overall signaling pathway. (d) Diagram of the *Spp1-Cd44* signaling network showing cell types signaling to neutrophils (left) and all other cell types (right). (e) Representative immunofluorescence microscopy for Neutrophil Gr-1 (red), OPN (green) and DAPI nuclear stain (blue) at 7 DPI left is a tilescan image, scale bar = 500 μm lesion site is to the left of the dotted white line, right is a zoomed in image below lesion site, scale bar = 50 μm (n = 3).

As neutrophils have not been widely examined as a source of OPN, we sought to identify the conditions that would induce neutrophil expression of this gene. We hypothesized that *Spp1* is not produced in BM or circulating neutrophils, but “environmental cues” upon reaching the infected tissue induce *Spp1* expression in neutrophils. To test this idea, we reanalyzed a publicly available dataset of neutrophils isolated from BM, blood and spleen from WT and E. coli infected mice [20]. Consistent with our hypothesis, we did not identify neutrophil subsets with high *Spp1* in the BM, blood, or spleen, even during infection (Supplemental Fig. 3a) To validate this observation in our infection model, we performed qPCR on neutrophils isolated from BM, blood, and skin lesions of *S. aureus* infected mice. We found that *Spp1* expression was only highly increased in neutrophils in the skin and not in blood or BM following skin infection (Supplemental Fig. 3b), supporting the idea that local tissue cues drive *Spp1* induction. To assess if this response was specific to *S. aureus* or a general feature of skin tissue inflammation, we performed a comparative transcriptomic analysis of our 7 DPI single cell dataset and a publicly available sterile mouse wound dataset [50] (Supplemental Fig. 3c). At 4 days post sterile wounding, we identified a skin neutrophil population producing high levels of *Spp1* (Supplemental Fig. 3d, 3e). These findings show that, following inflammation, neutrophils in the skin can express high levels of *Spp1* regardless of whether the stimulus was infectious or sterile. As both the sterile wound model and our dermonecrotic infection models are actively undergoing repair, it is conceivable that OPN may be playing a key functional role in promoting tissue regeneration during these processes.

### OPN deficiency reduces tissue repair cell types and signaling pathways

To investigate OPN’s role in tissue repair, we looked at early (7 DPI) and late (14 DPI) timepoints during tissue repair in our dermonecrotic skin infection model. We performed single cell transcriptomics on WT and OPN KO mice to evaluate cell type and signaling pathway changes during the tissue repair stage following skin infection.

Following read alignment and quality control filtering with Trailmaker software (Parse Biosciences), we performed comparative analyses of our samples using Seurat [51]. Figure 4a shows the reduced dimensionality clustering of cells in WT and OPN KO samples at the selected timepoints. After RPCA integration of all samples and clustering, we identified 13 major cell populations (Fig. 4b), including several types of immune cells, keratinocytes, and fibroblasts (Fig. 4c). Among the total keratinocytes, we performed subclustering and identified several keratinocyte subpopulations (Supplemental Fig. 4b, 4d). These included two stem cell populations: *Lgr5*+ HF stem cells and *Lgr6*+ HF stem cells (Supplemental Fig. 4d). HF stem cells have previously been shown to multiply and play a critical role in wound healing [52]. We also identified a population of epithelial cells that had similar gene expression to previously identified migrating wound edge epithelium (Mi_WE) [29] (Supplemental Fig. 4d). These included high levels of integrins, as well as *Krt16* and *Krt6a* (Supplemental Fig. 4e). In confirmation of this finding, we had observed KRT6A+ migrating cells along the lesion edge at 7 DPI (Fig. 1d).

**Figure 4.**
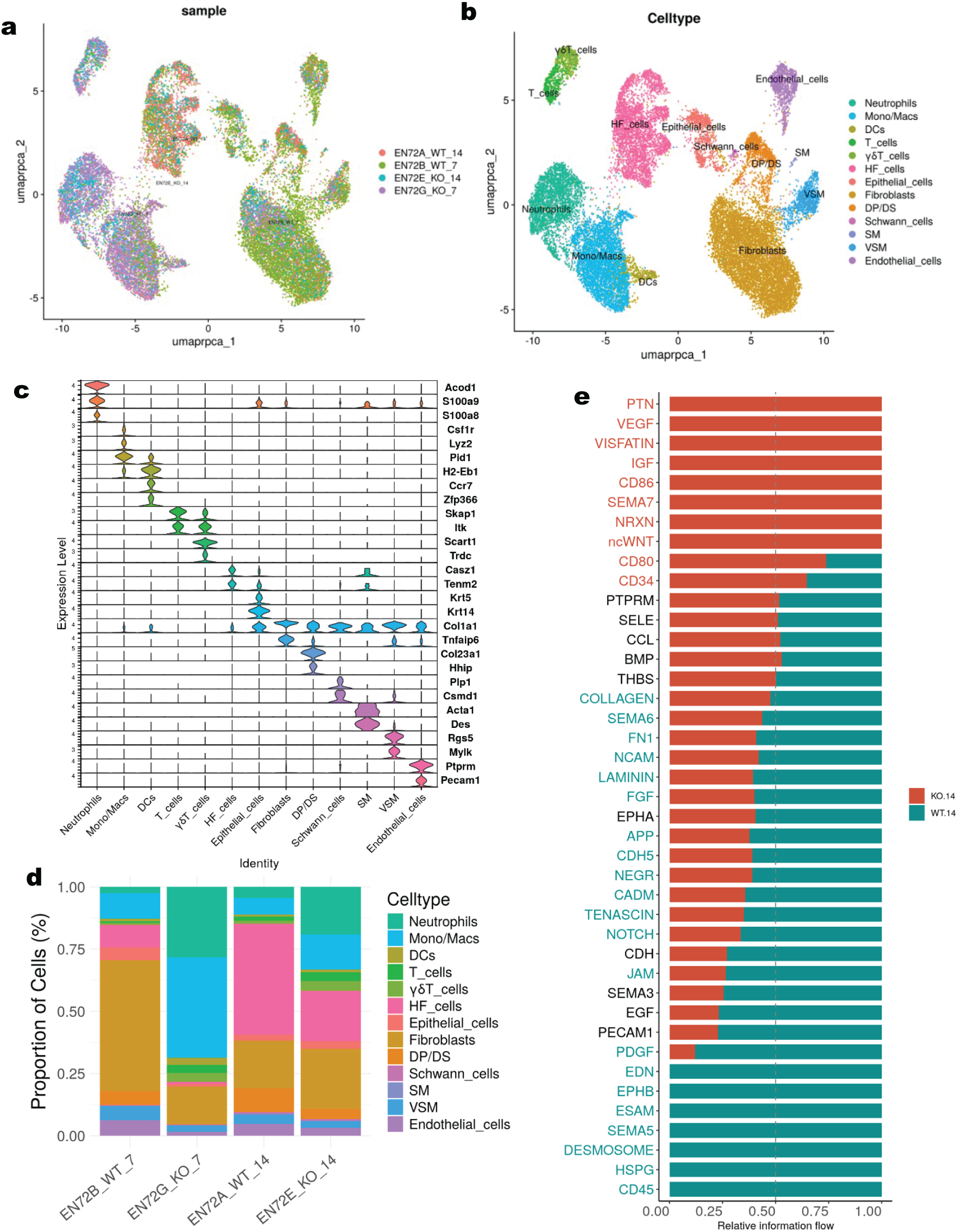
Comparative transcriptomic analysis of *S. aureus* infection dynamics in WT vs OPN KO mice. (a) UMAP dimensional reduction of WT and KO samples from the 7 DPI and 14 DPI timepoints (n = 3 mice per group, pooled). (b) UMAP of scRNA-seq skin cells showing 27 cell populations (c) Violin plots of representative genes used to identify cell types for each cell population identified in (b), left violin plot represents immune cells, middle plot is keratinocytes, and right plot is for fibroblast like and muscle cells. (d) Bar chart showing proportions of each cell type identified by scRNA-seq in KO and WT mice at each timepoint. (e) CellChat analysis of intercellular communication networks in KO (pink) and WT (blue) mice at 14 DPI. Pathways where the text is blue or pink represent pathways that are significantly upregulated in the WT or KO skin, respectively.

While the overall types of cells recruited in WT and OPN KO mice did not change, the relative proportions of several key repair cell types were markedly reduced in KO mice. At 14 DPI, WT mice exhibited a significantly higher proportion of fibroblasts, comprising approximately 58% of the lesion-associated cells, compared to only 15% in the OPN KO group (Fig. 4d). Additionally, keratinocyte (combined HF cell and epithelial cell) abundance was higher in WT mice at both 7 and 14 DPI. By 14 DPI, the proportion of total keratinocytes in WT mice was nearly double that observed in OPN KO mice (Fig. 4d; Supplemental Fig. 4c-d). These findings suggest that OPN plays a critical role in promoting fibroblast and keratinocyte recruitment or survival, potentially contributing to effective tissue repair following infection.

To further investigate signaling pathways differentially regulated between WT and OPN KO mice, we employed CellChat to analyze upregulated ligand–receptor interactions in each group. Several pathways known to be involved in tissue repair were significantly enriched in WT compared to OPN KO mice (Fig. 4e). These included DESMOSOME, HSPG, PDGF, NOTCH, and SEMA5. Collectively, these findings support our *in vivo* findings suggesting a critical role for OPN in promoting tissue repair during *S. aureus* skin infection by enhancing key regenerative signaling networks.

### OPN promotes lesion repair during *S. aureus* infection

Based on our scRNA-seq finding of reduced pro-repair cells and diminished regenerative signaling pathways in OPN KO compared to WT mice, we hypothesized that OPN KO mice would exhibit impaired tissue repair following *S. aureus* infection. To validate this hypothesis, we infected both OPN KO and WT mice following our dermonecrotic skin infection model and monitored external lesion size over time in both groups of mice (Fig. 5a). Consistent with our transcriptomic data, we found that OPN KO mice exhibited significantly larger skin lesions than WT controls (Fig. 5b) indicating an impaired healing response. To determine if the slower lesion resolution in OPN KO mice was due to impaired clearance of bacteria, we compared bacterial counts at 14 DPI. No significant differences in bacterial counts were observed between WT and OPN KO mice (Fig. 5c), suggesting that OPN does not influence bacterial clearance, but rather plays a role in lesion repair.

**Figure 5.**
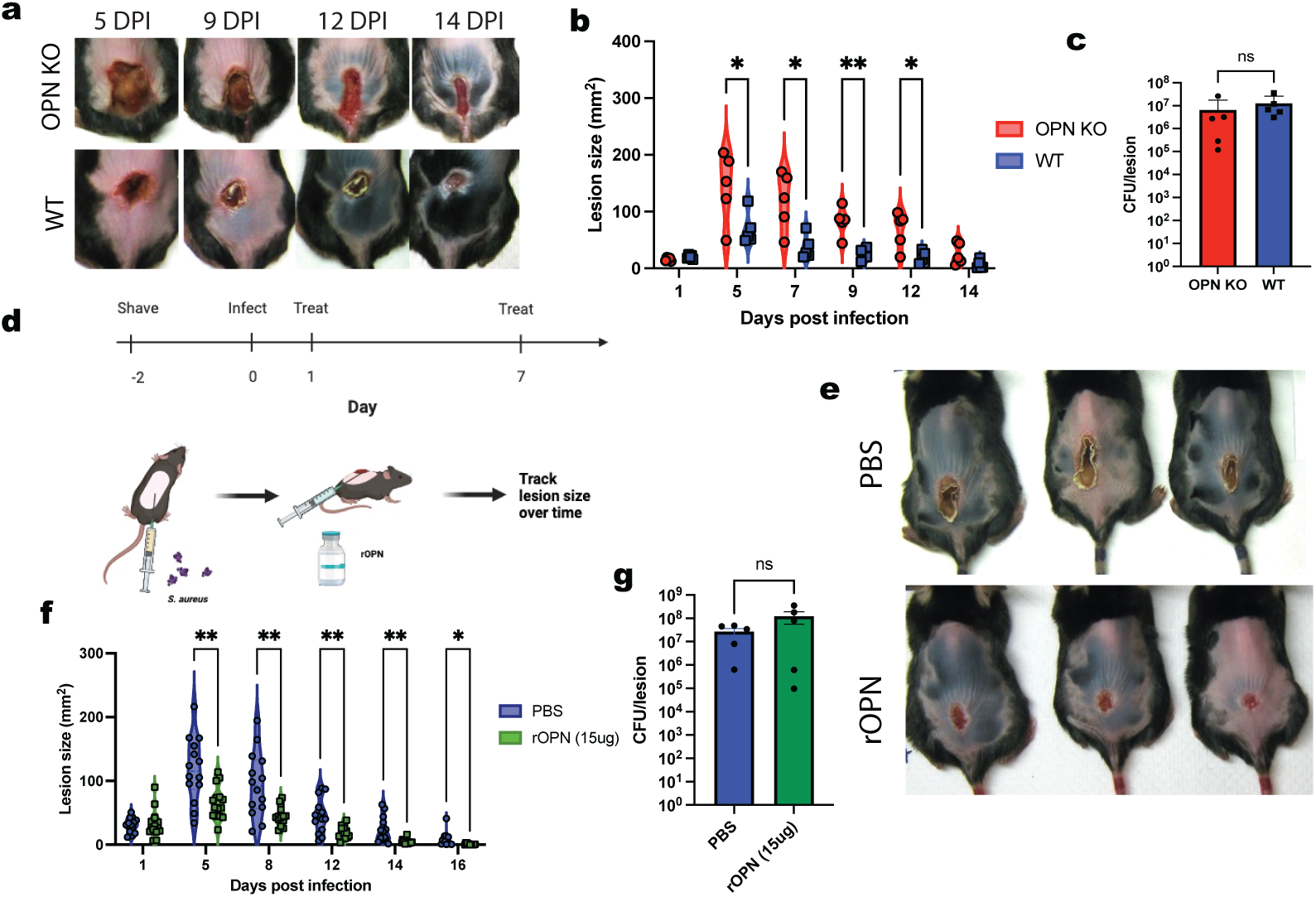
Osteopontin promotes lesion healing during *S. aureus* infection. (a) Representative images of skin lesions following infection in OPN KO and WT mice. (b) Skin lesion size during *S. aureus* infection in OPN KO (red) vs WT mice (blue) (n = 5 mice per group) (c) Bacterial load in the skin of WT and OPN KO mice 14 DPI (n = 5 mice per group). (d) Treatment schematic for rOPN and PBS. (e) Representative images of skin lesions from WT mice at 12 DPI, treated with rOPN (bottom) or PBS control (top). (f) Skin lesion size during *S. aureus* infection in WT mice treated with rOPN (green) or PBS control (blue) (n = 15 mice per group, at least 2 independent experiments). (g) Bacterial load in the skin of PBS (blue) and rOPN (green) treated WT mice at 14 DPI (n = 5 mice per group). Statistical analyses were performed using unpaired two-tailed Student’s *t* test. Means ± SEM, \**P* < 0.05, \*\**P* < 0.01, ns, not significant.

As we observed both transcriptomic changes and impaired tissue repair when we knocked out osteopontin, we investigated if treatment with recombinant osteopontin (rOPN) would enhance lesion repair during SSTI. To test this, WT mice received intradermal injections of rOPN (15 ug) or PBS control at the infection site once per week (Fig. 5d). Mice treated with rOPN had faster rounds of scabbing, with mice at 12 DPI still presenting with a scab in the control treated group whereas the rOPN treated mice had already shed their scab at the same timepoint (Fig. 5e). rOPN treated mice had significantly smaller lesions than PBS control treated mice starting at 5 DPI and experienced faster overall lesion resolution (Fig. 5f). Consistent with the knockout mouse model, treatment did not impact bacterial numbers present in the lesion (Fig. 5g). Taken together, these results show that OPN plays a critical role in tissue repair during skin infection and that therapeutic augmentation of OPN may be a promising strategy to improve healing outcomes.

## Discussion

In this study, combining transcriptomic and *in vivo* approaches, we identified a key role for Osteopontin (OPN) during skin infections. A major challenge in treating skin infections is delayed tissue repair and reepithelialization. Moreover, the host factors responsible for repair processes are poorly understood. To address this gap in knowledge, we performed scRNA-seq on infected mouse skin tissue and performed cell-cell communication analysis. From these studies, we discovered that OPN (SPP1) was the most upregulated signaling pathway during skin infection. The *Spp1* gene encodes a glycoprotein, OPN, which has broad ranging biological effects. Acting through the CD44 receptor and various integrins, secreted OPN promotes cell migration as well as longevity via anti-apoptosis functions [43, 53]. However, the role of OPN during skin infection has not been identified.

Remarkably, skin infection of OPN KO mice resulted in larger skin lesions than in WT mice despite no difference in bacterial clearance suggesting that OPN likely facilitates tissue repair. To further understand the repair processes that are OPN-dependent we compared single cell transcriptomics of OPN KO and WT mice after skin infection. Comparative analysis revealed large decreases in the proportion of fibroblasts and keratinocytes present at both 7- and 14-days post infection in OPN-KO mice. Fibroblasts play an important role in both matrix deposition and organization [54, 55], and keratinocytes including hair follicle stem cells are known to promote wound repair by expanding, differentiating and promoting reepithelialization[33, 52]. These decreases in repair associated cell types may contribute to the increase in lesion size observed in OPN KO compared to WT mice.

In addition to the differences in the types of cells present at the lesion site, we also found that several pathways with known roles in wound healing were downregulated in OPN KO mice. These included reductions in DESMOSOME, SEMA5, HSPG, PDGF and NOTCH signaling pathways. Desmosomes are critical mediators of cell-cell adhesion and the decrease of desmosomes in OPN’s absence may lead to decreased tissue integrity under stress [56]. Similarly, semaphorins, PDGF and HSPG signaling have all been shown to promote wound healing by increasing angiogenesis and epithelial cell migration following wounding [57–61]. PDGF also promotes deposition of extracellular matrix [62]. NOTCH signaling is important for the maintenance and proliferation of skin stem cells [63], which are critical during skin tissue repair [64]. Together, the decrease in these wound healing signaling pathways in OPN-KO mice could result in decreased barrier formation, keratinocyte migration, angiogenesis, and stem cell renewal, all potentially slowing tissue repair following skin infection.

As transcriptomic and *in vivo* data suggested involvement of OPN in infected tissue repair during skin infection, we tested if we could promote repair to speed lesion healing using an intradermal OPN injection and found that rOPN treatment sped tissue repair in WT mice following skin infection. These results underscore the potential to use rOPN as a therapeutic treatment to improve tissue repair during skin infection. As rOPN did not have an impact on bacterial load, it could conceivably be given in conjunction with antibiotics to improve overall therapeutic outcomes.

In addition to the role of OPN and its therapeutic value during infection, our study addresses the important question of the source of OPN in the infected skin. Prior to our studies, neutrophils were not thought of as major producers of *Spp1* [48]. While we did see some *Spp1* expression from the mono/mac population, as expected, a population of neutrophils were unexpectedly the largest producer of *Spp1* in the lesion site of our model. While neutrophils are known for their role in the containment and killing of bacteria [65], recent work on neutrophil diversity has suggested that neutrophils are not homogenous and different neutrophil subtypes could play roles beyond bacterial clearance [20, 66]. It has also been shown that neutrophils can experience reprogramming once they enter different tissues [21]. Our findings that high *Spp1* is not present in BM or blood neutrophils but only skin neutrophils provide key insights into the origin of the *Spp1*^high^ neutrophil subset. Based on these findings, it is likely that some of the neutrophils entering the skin experience distinct transcriptional changes to enter a pro-repair state and turn on expression of *Spp1*.

In addition to the *Spp1*^high^ Neut4 subtype, through transcriptomics, we identified multiple other neutrophil populations present during skin infection. This heterogeneity could be a mechanism to allow neutrophils to undertake multiple roles in the infection site to facilitate both classical bacterial clearance as well as tissue repair functions simultaneously. We identified a classical bacterial killing subtype expressing TNF, Complement, and CXCL signaling. We also found neutrophil subtypes with upregulation of gene pathways associated with tissue repair including GALECTIN, CD80, and ANNEXIN. Annexin signaling is involved in resolving inflammation in the skin and in promoting collagen deposition by fibroblasts [67]. Treatment with AnxA1 peptides have also been shown to improve wound healing [68, 69]. *Lgals9*, a galectin gene, has been shown to stimulate keratinocyte proliferation when IL-17 is present [37], like during *S. aureus* infections. Together, these neutrophils may promote tissue repair through boosting fibroblast activity and promoting keratinocyte expansion. In addition to tissue repair, coordination of the cells via cell-cell signaling is a critical process during infection response. The *Spp1*^high^ neutrophil group had upregulated GO terms related to chemokine and cytokine activity as well as Toll-like receptor signaling, C-type lectin receptor signaling, and NF-κB signaling, indicating that these neutrophils may play a key cell-cell signaling role. We also identified a subtype likely involved in antigen presentation based on enrichment of CD80 signaling and transporter associated with antigen processing (TAP) binding genes which are involved in MHC presentation [70]. While antigen presentation is not a classical role for neutrophils, ex vivo studies have shown that neutrophils pulsed with antigen elicit robust IL-17 responses from T cells [71]. IL-17 is a critical cytokine for the resolution of *S. aureus* skin infections [10, 11]. The antigen-presenting neutrophil subtype that we observed during infection may play a role in skin infection resolution by promoting the IL-17 γδ T cells response. Neutrophils were the predominant cell type within our infected tissue; the studies we conducted have shown multiple different roles for neutrophil subsets to allow for the coordinated resolution of *S. aureus* skin infection.

Altogether, this study has advanced our understanding of the tissue repair process during skin infection. Including identifying important roles for OPN and heterogeneous neutrophil subtypes in this process. Importantly, we were able to translate this discovery into a therapeutic strategy, using rOPN, that significantly improved tissue repair during skin infection in WT mice. In future, this therapy could potentially be used in conjunction with antibiotics to improve outcomes for patients with skin infections.

## Acknowledgements

We would like to thank Jessica Shannon from J. Zhang’s laboratory (Duke University) for technical instruction of skin processing, sectioning and staining. We thank Wanying Miao and Yingai Jin from the Zhang lab for single cell dissociation protocol development. We thank the Duke University School of Medicine for the use of the Sequencing and Genomic Technologies Shared Resource, which provided sequencing and basic data analysis services. We would like to acknowledge the assistance of the Duke Molecular Physiology Institute Molecular Genomics Core for the generation of scRNA-seq data for the manuscript. We would like to thank the Duke Immunology Flow Cytometry Core for access to their facilities. Figure schematics were created with <u>BioRender.com</u>.

## Author Contributions

Studies were designed by E.E.N, A.P.S.R, M.L.S, J.Y.Z, and S.N.A. Experiments were performed by E.E.N, and experimental data were analyzed by E.E.N and A.P.S.R with advice from M.L.S, J.Y.Z, and S.N.A. Sequencing data were analyzed by E.E.N, with preliminary alignment and filtering performed by Duke Molecular Genomics Core. The manuscript was primarily written by E.E.N, A.P.S.R, and S.N.A. All authors contributed to paper discussions and review of the manuscript.

## Materials and Methods

### Mouse strains

For this study, female C57BL/6J (stock no. 000664) and OPN KO (stock no. 004936) mice were purchased from Jackson Laboratories. Mice used for experiments were eight to ten weeks of age.

Mice were housed in Duke University–approved facilities and received water and food ad libitum. Experiments were performed in accordance with a Duke University Institutional Animal Care and Use Committee–approved animal protocols.

### Dermonecrotic skin infection model

Skin infections were based on a dermonecrotic infection model that we have previously described [24, 72]. All skin infections from this study were performed with clinical strain *S. aureus* (strain ID 10201) grown in Tryptic Soy Broth (BS Biosciences). Bacteria were grown overnight, shaking at 37C before being diluted 1:1000 and incubated until they reached log phase. Two days prior to infection, mice had their dorsal hair removed using clippers followed by application of hair removal cream (Veet). Mice were injected intradermally in the middle of the shaved dorsal region with a 100 ul dose containing 5E7 CFU log-phase bacteria complexed with dextran microbeads (50g/L, Cytodex, Sigma-Aldrich) in phosphate buffered saline (PBS). The microbeads were used, as previously described, to create a localized skin infection and prevent bacterial dissemination. Lesion sizes were assessed by digital planimetry, where lesion outlines were traced on acetate transparency films (Staples) and ImageJ were used to calculate surface areas of the lesion tracings.

### Colony forming unit (CFU) assay

To determine bacterial burden, infected skin tissues were harvested at different points post infection and homogenized in cold 0.1% Triton X-100 in PBS for 3 cycles of 90s with 10 minutes on ice between cycles using an automatic homogenizer. Serial dilutions were plated in duplicates on TSB agar plates, and colonies were counted after overnight incubation at 37°C.

### Intradermal treatments

Carrier free recombinant mouse osteopontin (rOPN, R&D Systems) was dissolved in sterile PBS. Mice were treated at 1 and 8 DPI with either 15 µg of rOPN or PBS only control in a total volume of 50ul injected intradermally at the lesion site.

### Skin dissociation and single cell isolation

Single cell isolation was modified from previously described skin dissociation methods [73]. Briefly, lesions were collected, cut into small pieces, and incubated in a 37C water bath for 30 minutes in 300 µg/ml of Liberase TM (Roche) and 50 µg/ml DNAse I (Sigma) in RPMI (Thermo Fisher Scientific). Cells were spun down and resuspended in 0.05% Trypsin (Thermo Fisher Scientific), then incubated for 10 minutes at 37C, prior to being passed through a 70 µm filter into a new conical with RPMI + 5% FBS. Filters were placed in a 6 well plate, covered with 0.05% trypsin and incubated for 10 minutes at 37C before being placed back on conical tubes with corresponding cells and filters were washed with RPMI + 5% FBS to detach remaining cells. Cells were washed with cold 1% BSA in PBS for scRNA-seq, or 3% FBS and 5mM EDTA in PBS for flow cytometry.

### Flow Cytometry

After isolation of single cells, washed cells were blocked in 3% FBS and 5 mM EDTA in PBS buffer with 1% anti CD16/CD32, 5% normal rat serum, and 5% normal mouse serum. Cells were then stained with fluorochrome-tagged antibodies for surface markers. For any experiments involving intracellular staining, cells were fixed and permeabilized for intracellular staining using the BD Cytofix/Cytoperm kit (BD Biosciences) in accordance with manufacturer instructions prior to intracellular staining. Data were collected using a LSRFortessa X-20 Cell Analyzer (BD Biosciences) prior to analysis with FlowJo software (TreeStar).

### scRNA-seq

#### 10X Genomics

Following single cell isolation of cells from uninfected and 7 DPI skin from WT mice, dead cells were removed using the MojoSort Mouse Dead Cell Removal Kit (Biolegend) in accordance with manufacturer’s protocols. Cells were then submitted to the Duke Molecular Genomics Core for library preparation using the 10X Genomics Chromium Single Cell 3′ GEM kit and, Library and Gel Bead Kit v3. Libraries then were submitted for Illumina NovaSeq6000 SP sequencing and basic data analysis to the Duke Center for Genomic and Computational Biology. For data analysis, raw base call (BCL) files were demultiplexed into FASTQ files and aligned to the mm10 mouse reference genome, performed filtering, barcode and UMI counting was then done using 10X’s Cell Ranger software.

#### Parse

Following single cell isolation of cells from OPN KO and WT mice at 7 DPI and 14 DPI, washed cells were blocked with 1% anti CD16/CD32 with 5% normal rat serum, and 5% normal mouse serum in 1% BSA. Cells were then stained with fluorochrome-tagged antibodies for Ly6G (Biolegend) and viability marker 7-AAD (BD Biosciences). Ly6G+ and Ly6G- live cell populations were then sorted via fluorescent activated cell sorting (FACS) and mixed at a 50/50 ratio. Cells were then fixed using the Evercode Cell Fixation v3 kit (Parse Biosciences) in accordance with manufacturer’s protocols and stored at -80C before library preparation. Library preparation and barcoding were performed using the Evercode WT v3 kit in accordance with manufacturer’s protocol. Libraries were submitted to the Duke Center for Genomic and Computational Biology for sequencing using Illumina NovaSeq X Plus. For data analysis, raw base call (BCL) files were demultiplexed into FASTQ files which were then loaded into Parse Bioscience’s Trailmaker pipeline for alignment and filtering.

#### Downstream analysis

Downstream analyses were performed with Seurat (version 5) [51]. We filtered out cells with less than 200, or more than 2500 genes, and more than 7% mitochondrial reads. Then performed data normalization, scaling and principal component analysis (PCA). Samples from each batch were integrated using RPCA integration [51] followed by finding neighbors, clusters and UMAP dimensional reduction. Cell annotation was done based on differential gene expression from identified first-level clusters. In Parse dataset, keratinocytes were subclustered using Seurat and cell types were annotated based on second-level clustering. Secondary analysis to assess cell-cell communication was done using CellChat [42]. Signaling pathways expressed in less than 10 cells were excluded from final analysis.

#### Publicly available scRNA-seq datasets

Data from Xie et al. [20] (GSE137539) and Vu et al. [50] (GSE188432) were downloaded from NIH GEO and reanalyzed in accordance with the above protocol. Following clustering and UMAP reduction, neutrophils were identified and expression levels of *Spp1* were examined in the neutrophil populations.

### RNA isolation and qPCR

Skin lesions were cut into small pieces then homogenized in Trizol buffer (Thermo Fisher Scientific) using an automatic homogenizer for 3 cycles of 60s with 10 min on ice between cycles. RNA was isolated in accordance with manufacturer’s protocols. RNA purity and concentration was assessed using a Nanodrop. RNA was reverse transcribed using the iScript cDNA Synthesis Kit (Biorad). Quantitative PCR was performed using iQ SYBR Green Supermix (BioRad) with the following primers: *Il1b*-forward 5’-TTGACGGACCCCAAAAGAT-3’, *Il1b*-reverse 5’-GAAGCTGGATGCTCTCATCTG-3’, *Gapdh*-forward 5’-TGACCTCAACTACATGGTCTACA-3’,

*Gapdh*-reverse 5’-CTTCCCATTCTCGGCCTTG-3’, *Spp1*-forward 5’-AGCAAGAAACTCTTCCAAGCAA-3’, *Spp1*-reverse 5’-GTGAGATTCGTCAGATTCATCCG-3’ (Integrated DNA Technologies). qPCR was performed using the QuantStudio 6 Pro (Thermo Fisher Scientific). Relative gene expression was quantified using *Gapdh* expression in the same cDNA as internal control using the comparative Ct method.

### Scanning Electron Microscopy (SEM)

Skin lesions from mice at day 7 post infection were collected and fixed overnight in 4% PFA at 4°C. The tissues were dehydrated in an ethanol series before being dried in hexamethyldisilane (Sigma Aldrich, 440191). Lesion samples were sputter coated for 600 s with gold using the SPUT6 Vacuum Sputter Coater (Kurt Lesker PVD 75). Skin lesions were then imaged with a scanning electron microscope (FEI XL30 SEM-FEG).

### Immunofluorescence imaging

Skin from the lesion site was dissected and fixed in 4% PFA for 12 to 24 hours, followed by a sucrose gradient of 15% sucrose and 30% sucrose in PBS for 24 to 48 hours at 4C. Skin was then embedded in optimal cutting temperature (OCT) compound then frozen and stored at -80C until sectioning. 15-20 µm sections were fixed using 4% PFA and blocked using 2.5% normal donkey serum, 1% BSA, 1% Gelatin, 0.1% triton X-100 in PBS. Sections were stained with anti-mouse primary antibodies overnight at 4C in blocking solution. All sections were washed 3 x 10 mins in 0.01% triton X-100 in PBS, then stained with Donkey-anti secondaries to the species of the primary antibody 1:1000 in blocking buffer for 1 hour at RT. Sections were washed 3 x 10 mins in 0.01% triton X-100 in PBS and mounted using Fluoroshield with DAPI.

## Supplementary Figures

**figure S1.**
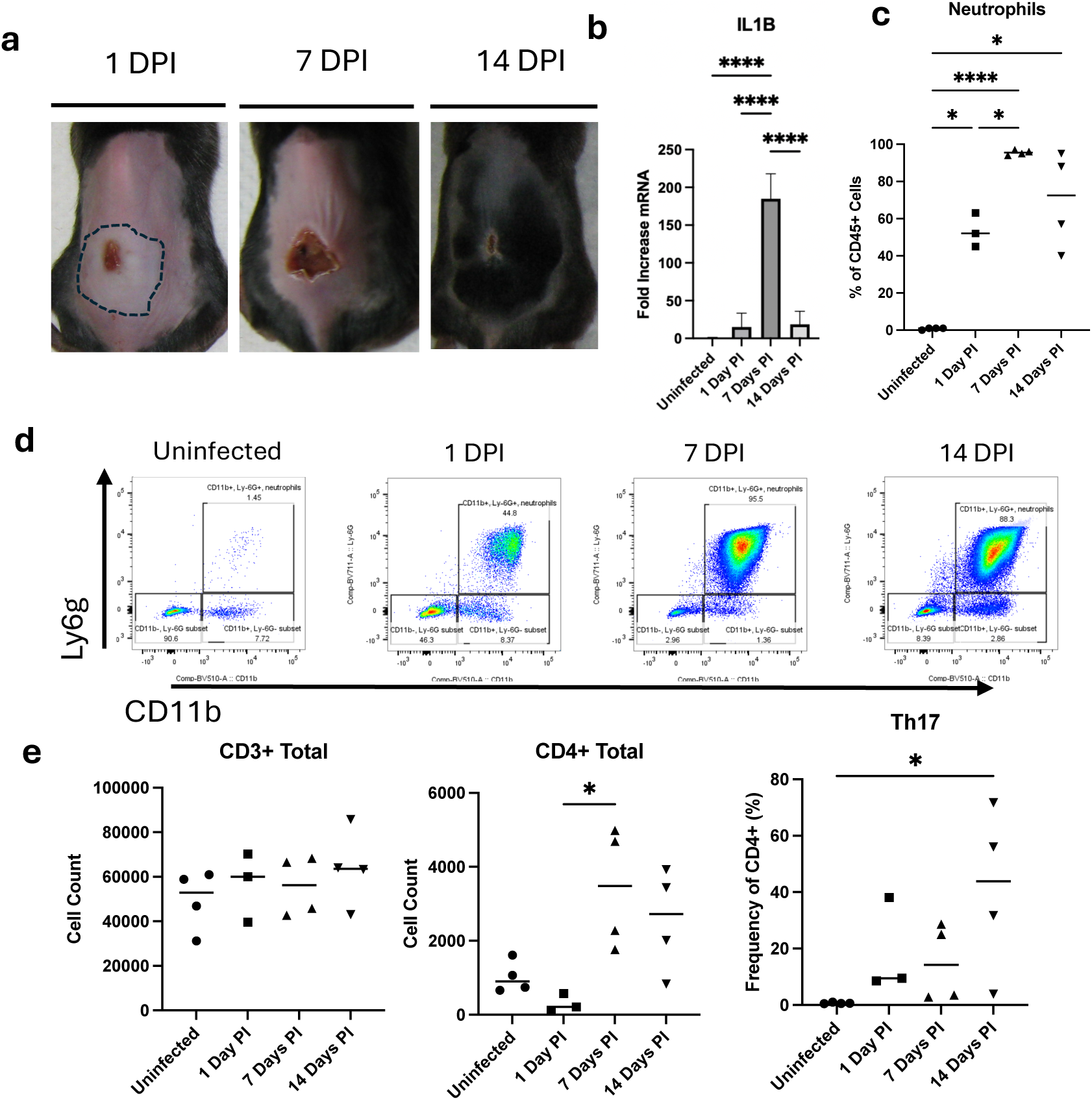
Skin lesion progression and immune response. (a) Representative images of dermonecrotic infection lesions over time. At 1 DPI there is a closed lesion where total inflamed area is outlined by black dashed line, followed by scabbing and start of closure at 7 DPI and progressive wound closure at 14 DPI. (b) Relative mRNA expression of *Il1b* over the course of infection, *Gapdh* was used as an internal control. (c) Flow cytometry quantification of percentage of neutrophils out of the total CD45+ immune cell population present at the skin lesion site at indicated timepoints post infection (d) Representative flow cytometry plots depicting the CD11b+Ly6G+ neutrophil populations at the skin lesion at di;erent timepoints. Cells shown were initially gated for size by FSC and SSC and single cells, followed by 7AAD-CD45+ gating for live immune cells. (d) Flow cytometry quantification of adaptive immune cells present in the skin lesion at indicated timepoints post infection. \**P* < 0.05, \*\**P* < 0.01, \*\*\**P* < 0.001, \*\*\*\**P* < 0.0001. PI, post infection.

**figure S2.**
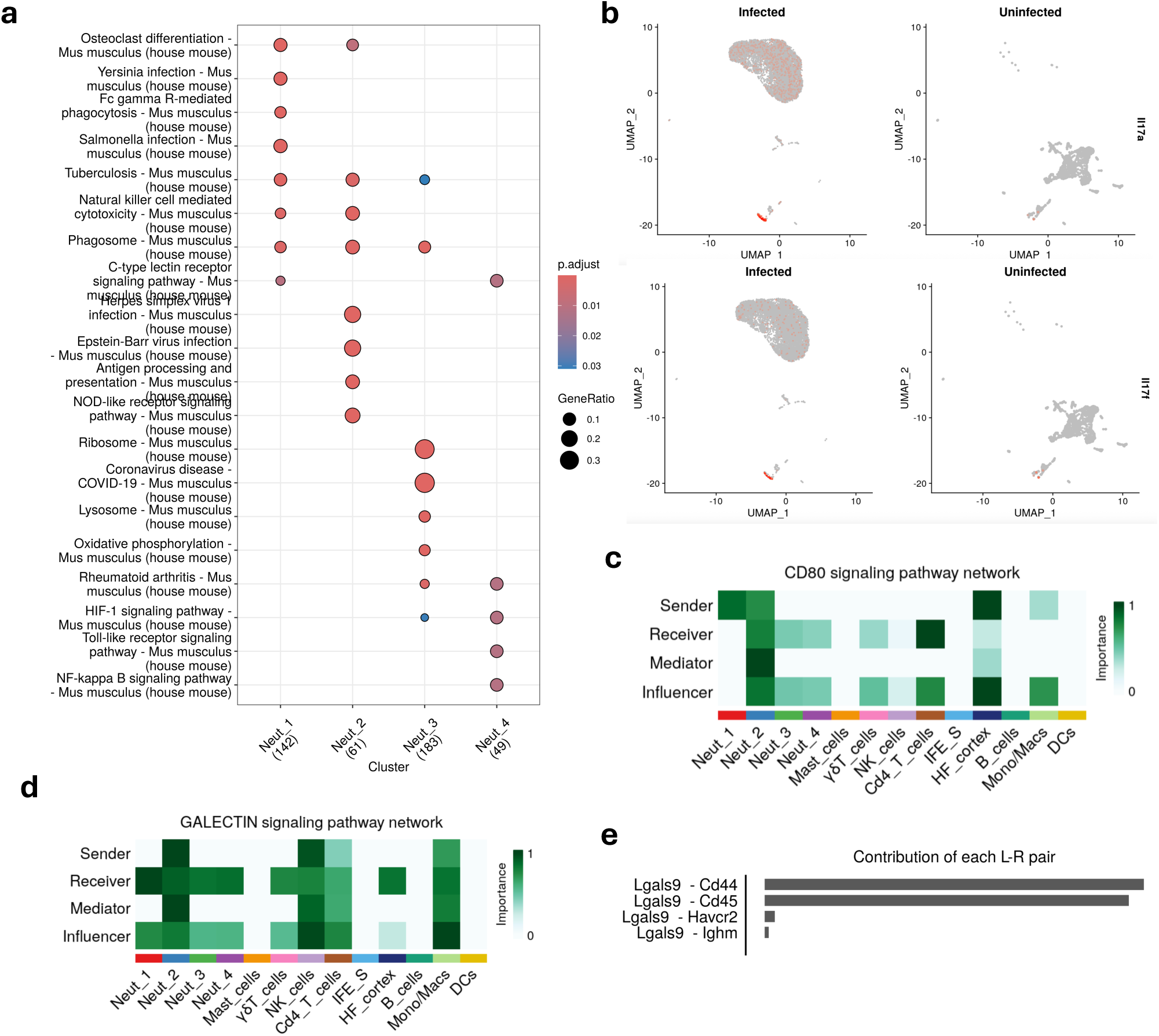
ScRNA-seq signaling responses during infection. (a) KEGG enrichment analysis of upregulated activity for neutrophil subpopulations identified via scRNA-seq at 7 DPI in mouse skin lesion. (b) UMAP plot of *Il17a* and *Il17f* expression by cells in infected skin lesion at 7 DPI vs in uninfected skin. Orange color represents cells with increased gene expression. (c) Cellchat heatmap of CD80 signaling pathway network for cell types from infected skin at 7 DPI. (d) Cellchat heatmap of Galectin signaling pathway network for cell types from infected skin at 7 DPI. (e) Cellchat heatmap of CD80 signaling pathway network for cell types from infected skin at 7 DPI. (f) Contribution of each ligand receptor pair in the GALECTIN signaling network to the overall signaling pathway at 7 DPI.

**figure S3.**
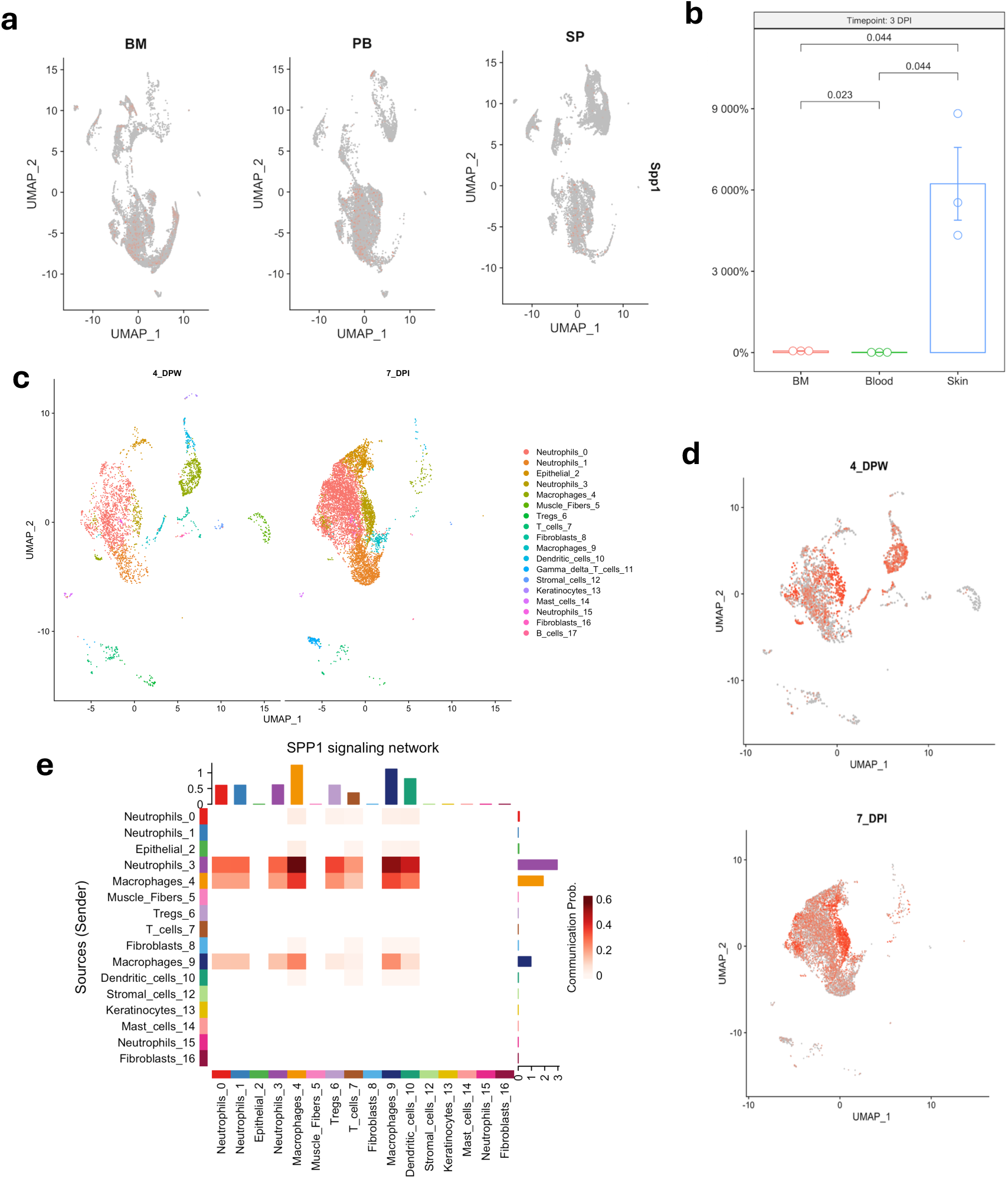
Comparative analysis of Spp1 in diUerent tissues and conditions. (a) UMAP plots from reanalysis of Xie et al existing dataset showing Spp1 expression levels in neutrophils isolated from bone marrow (BM), peripheral blood (PB), and spleen (SP). (b) Relative mRNA expression of *Spp1* in neutrophils isolated from the BM, blood, and skin following dermonecrotic *S. aureus* skin infection, GAPDH was used as an internal control. (c) UMAP showing cell types from comparative analysis of our 7 DPI scRNA-seq data and existing scRNA-seq dataset of skin at 4 days post wounding (DPW) from Vu et. Al. (d) UMAP plots showing *Spp1* expression levels in infected vs wounded skin from analysis in (c). (e) Cellchat heatmap of *Spp1* signaling senders and receivers present at 4 days post wounding.

**figure S4.**
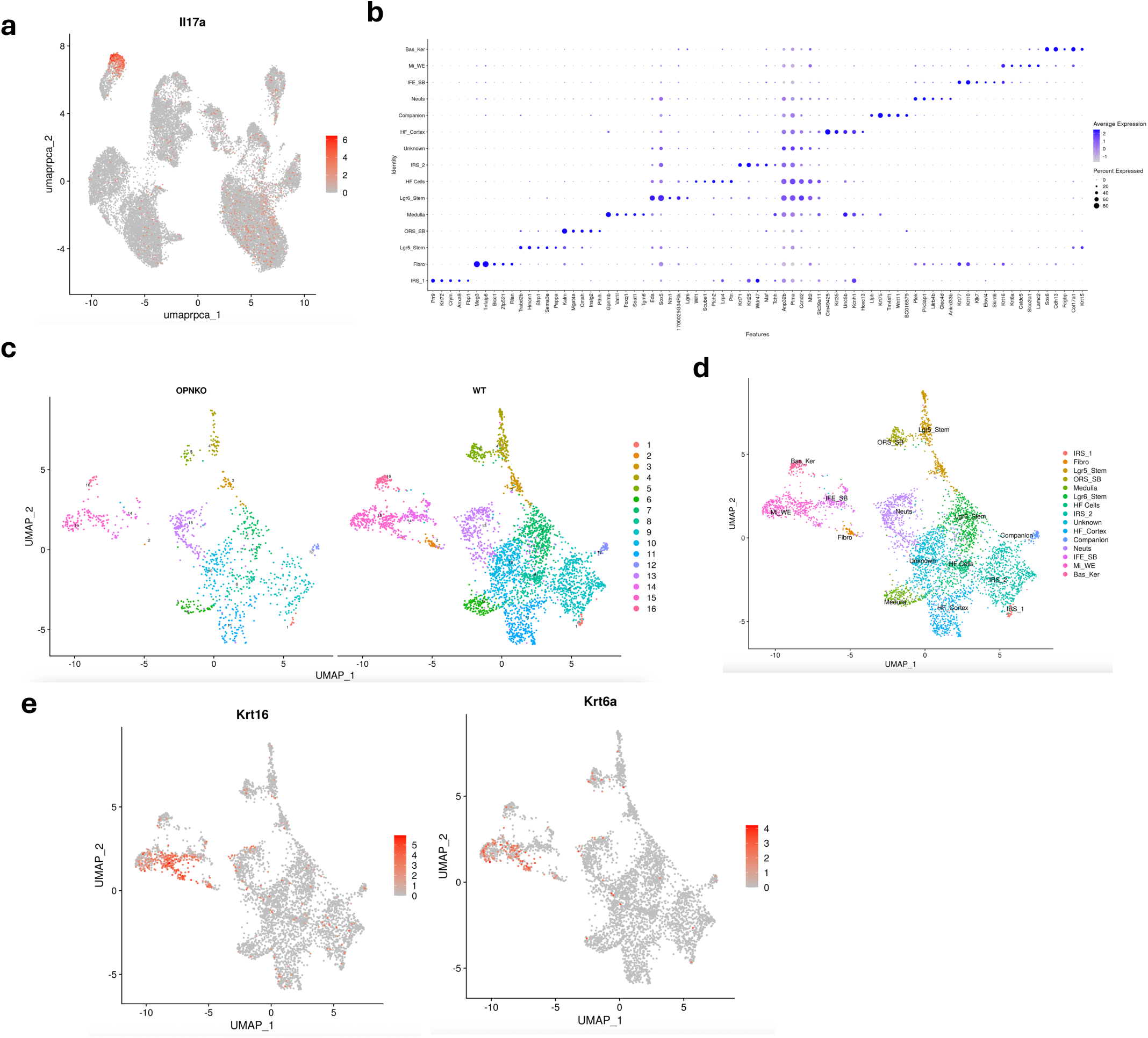
OPN KO and WT gene expression and keratinocyte subclustering. (a) UMAP *IL17a* expression levels are high in gd T cells (b) Dotplot of representative genes used to identify cell types following subclustering of keratinocytes from OPN KO and WT mice scRNA-seq data. (c) UMAP of scRNA seq subclustering of keratinocytes shows cluster distribution in OPN KO vs WT mice(d) UMAP plot showing identified cell types from keratinocyte subclustering based on genes in (b). (e) UMAPs of *Krt16* and *Krt6b* expression in keratinocyte subtypes show high levels in migrating wound edge (Mi_WE) population from (d).

## Notes

### Competing Interest Statement

The authors have declared no competing interest.

