## Supplemental Figures for "Osteopontin promotes lesion repair during *Staphylococcus aureus* skin infections"

**Supplementary Figures**

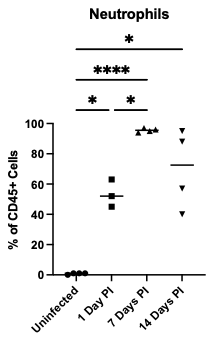

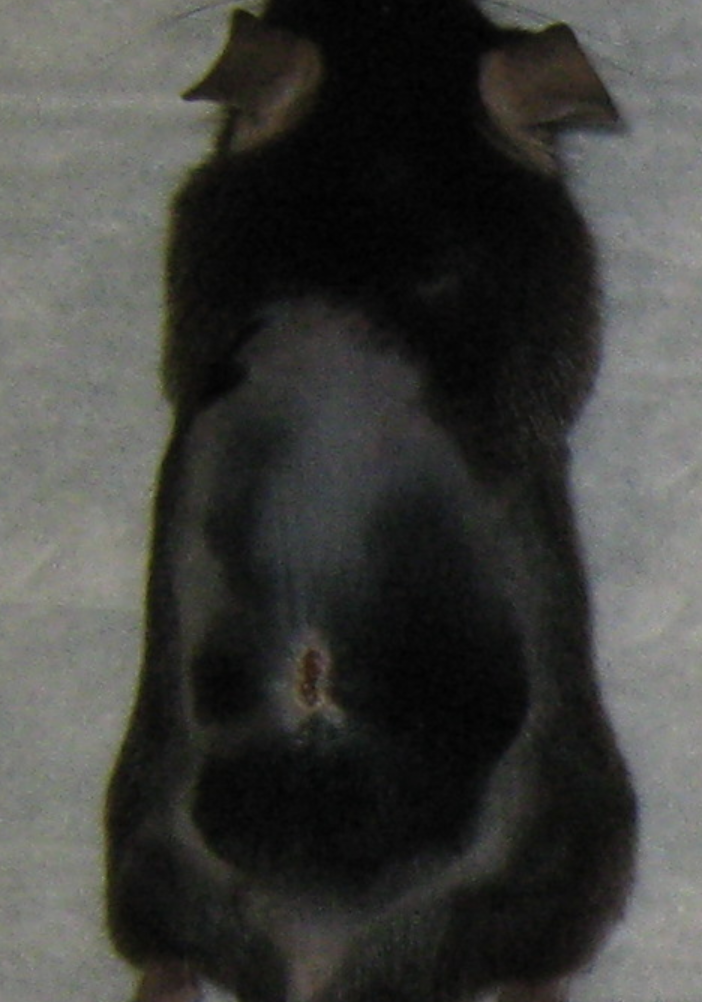

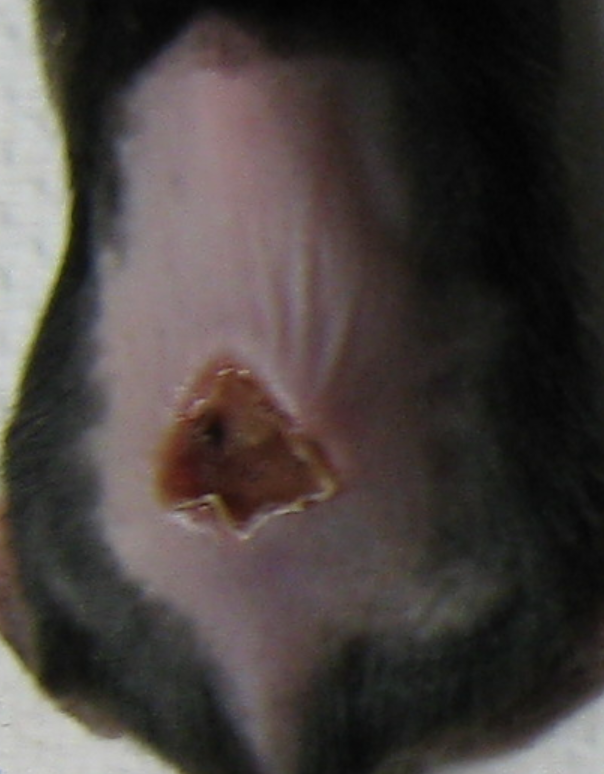

1 DPI

7 DPI

14 DPI

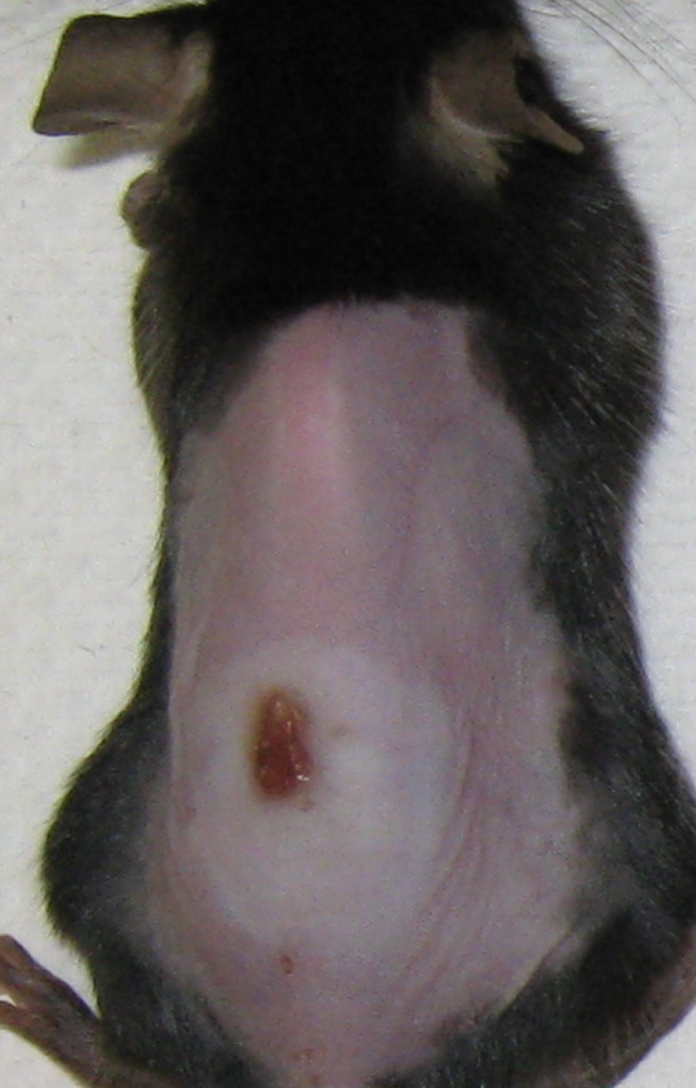

**a**

**b**

**c**

**e**

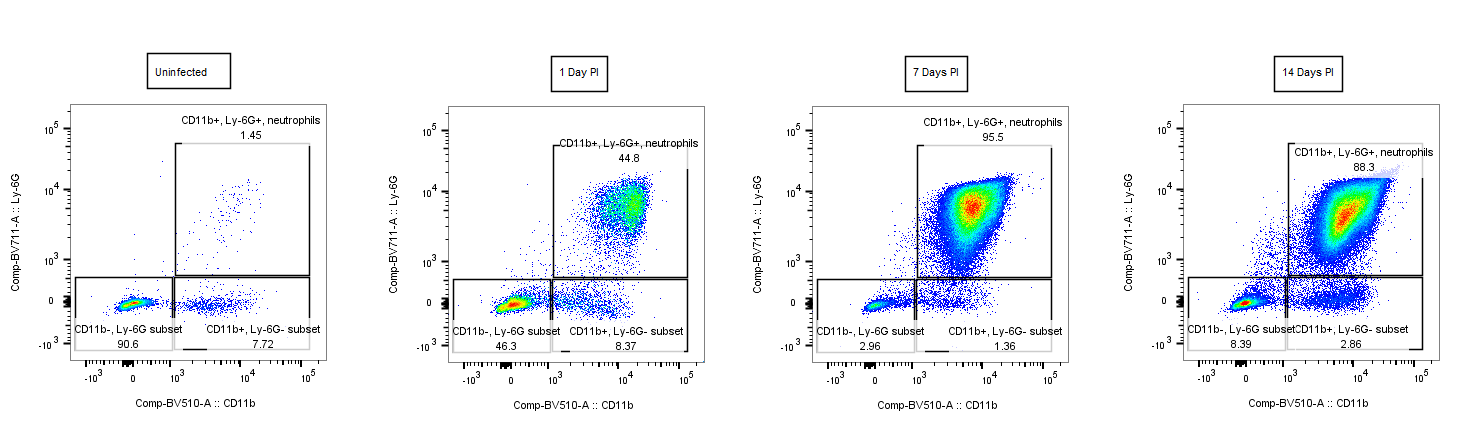

Uninfected

1 DPI

7 DPI

14 DPI

Ly6g

CD11b

**d**

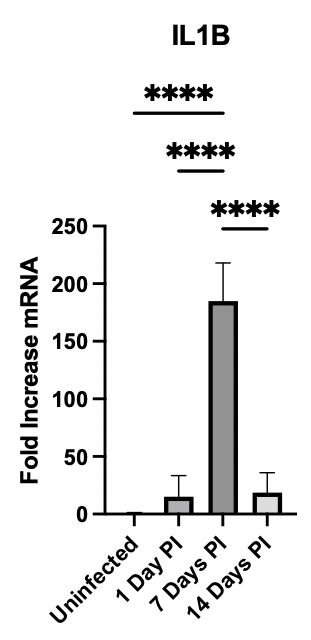

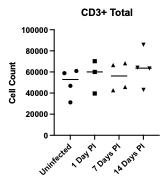

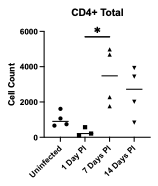

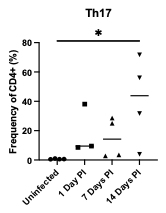

**figure S1. Skin lesion progression and immune response**. (a) Representative images of dermonecrotic infection lesions over time. At 1 DPI there is a closed lesion where total inflamed area is outlined by black dashed line, followed by scabbing and start of closure at 7 DPI and progressive wound closure at 14 DPI. (b) Relative mRNA expression of *Il1b* over the course of infection, *Gapdh* was used as an internal control. (c) Flow cytometry quantification of percentage of neutrophils out of the total CD45+ immune cell population present at the skin lesion site at indicated timepoints post infection (d) Representative flow cytometry plots depicting the CD11b+Ly6G+ neutrophil populations at the skin lesion at different timepoints. Cells shown were initially gated for size by FSC and SSC and single cells, followed by 7AAD-CD45+ gating for live immune cells. (e) Flow cytometry quantification of adaptive immune cells present in the skin lesion at indicated timepoints post infection. **P* < 0.05, ***P* < 0.01, ****P* < 0.001, *****P* < 0.0001. PI, post infection.

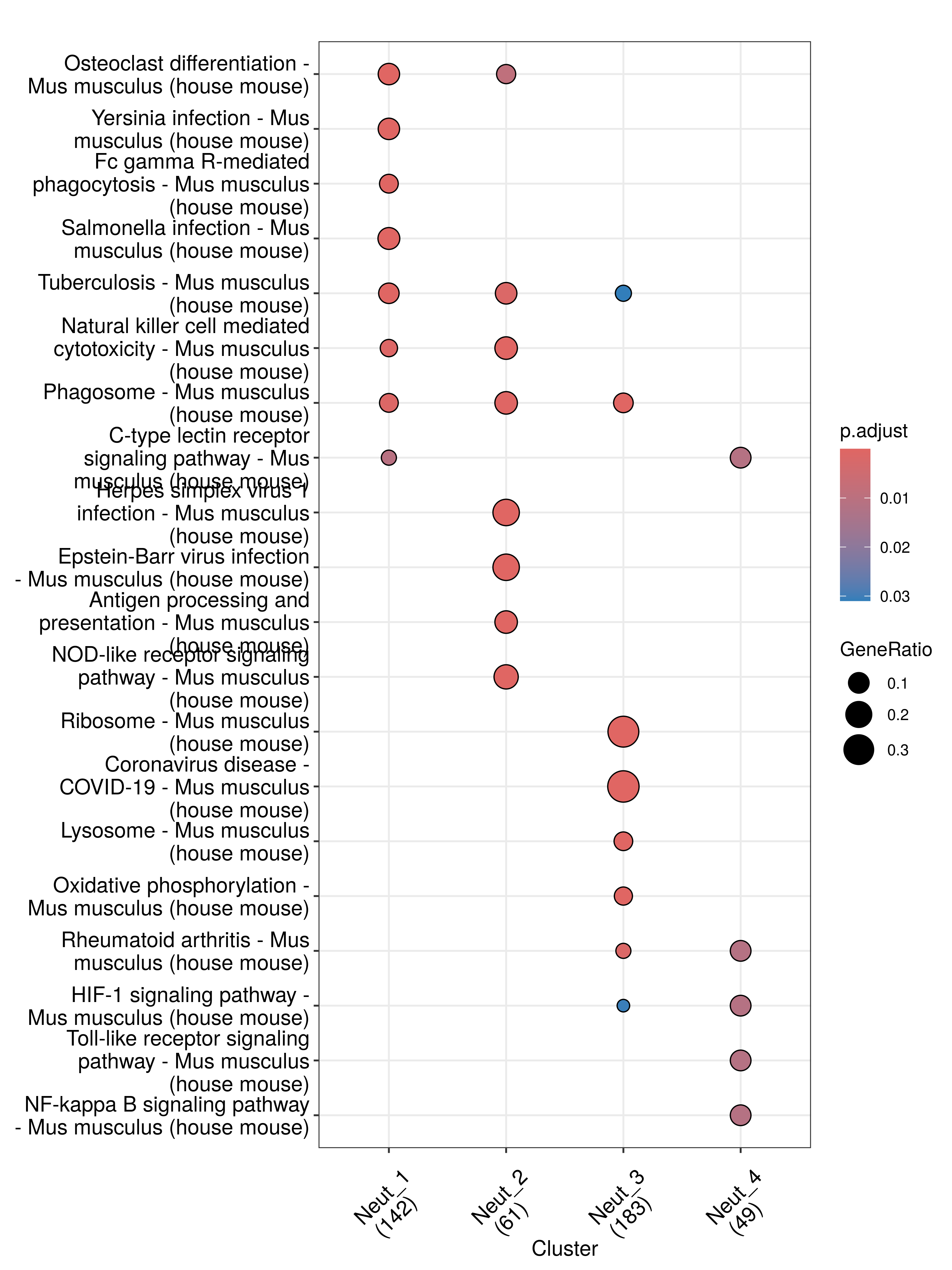

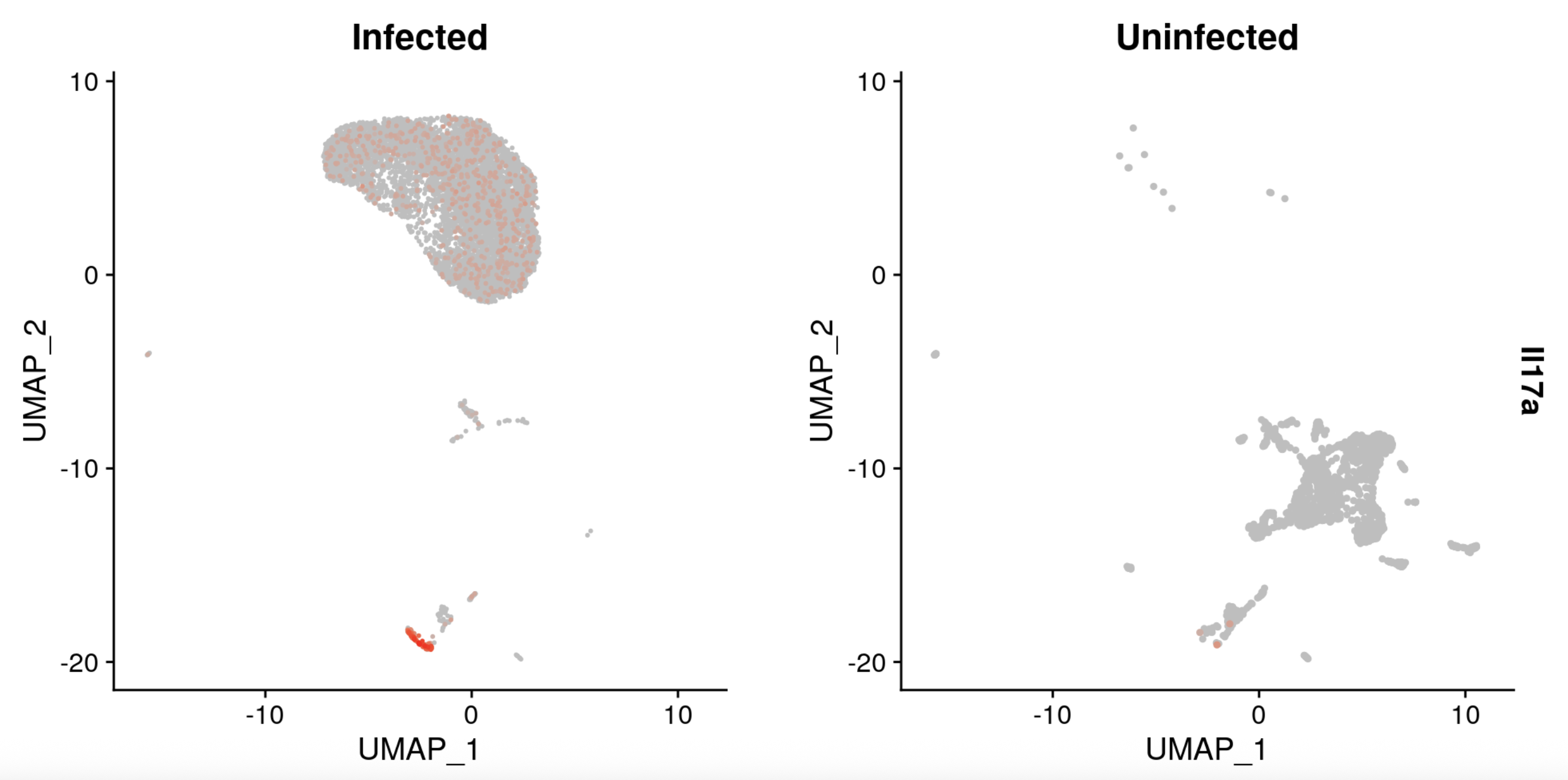

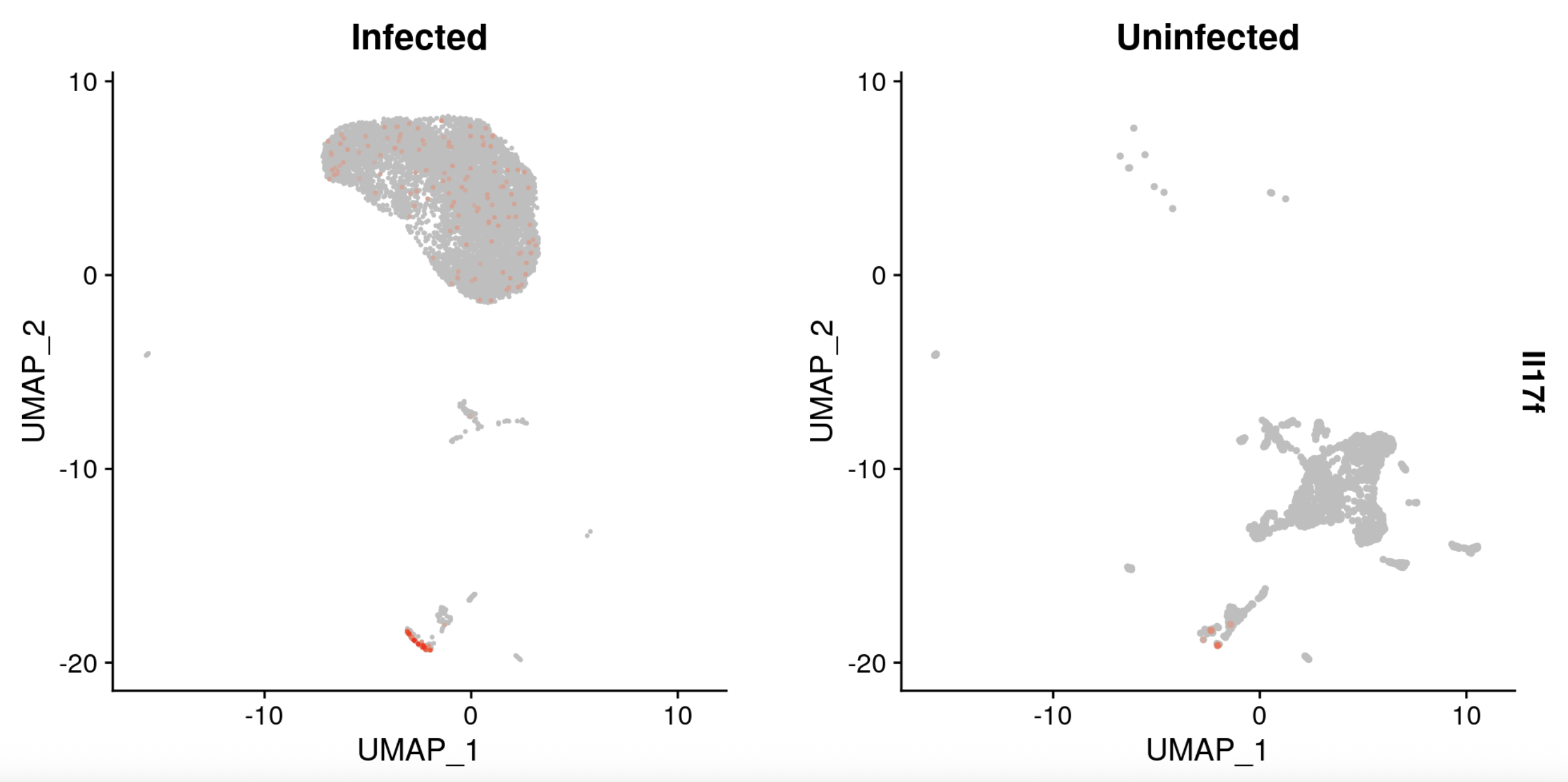

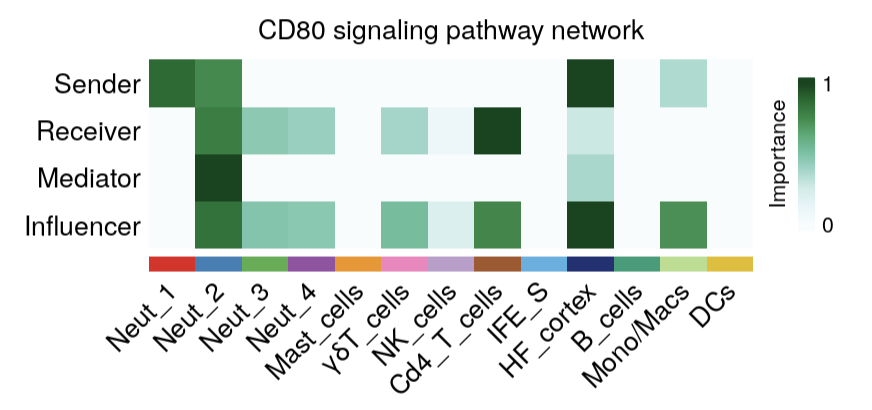

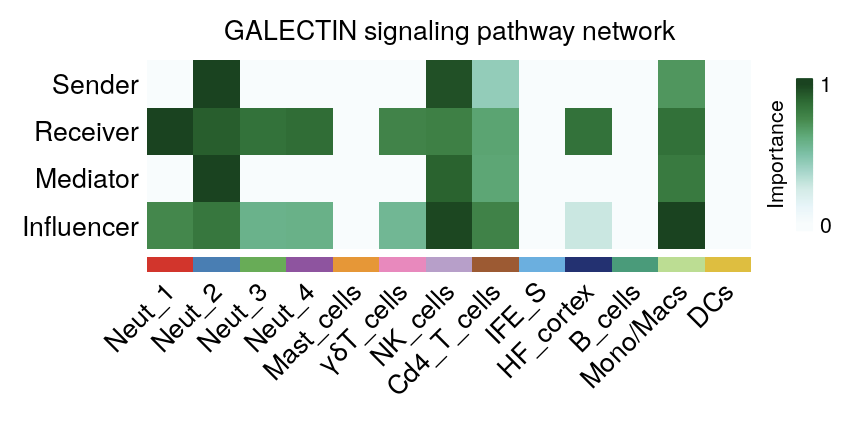

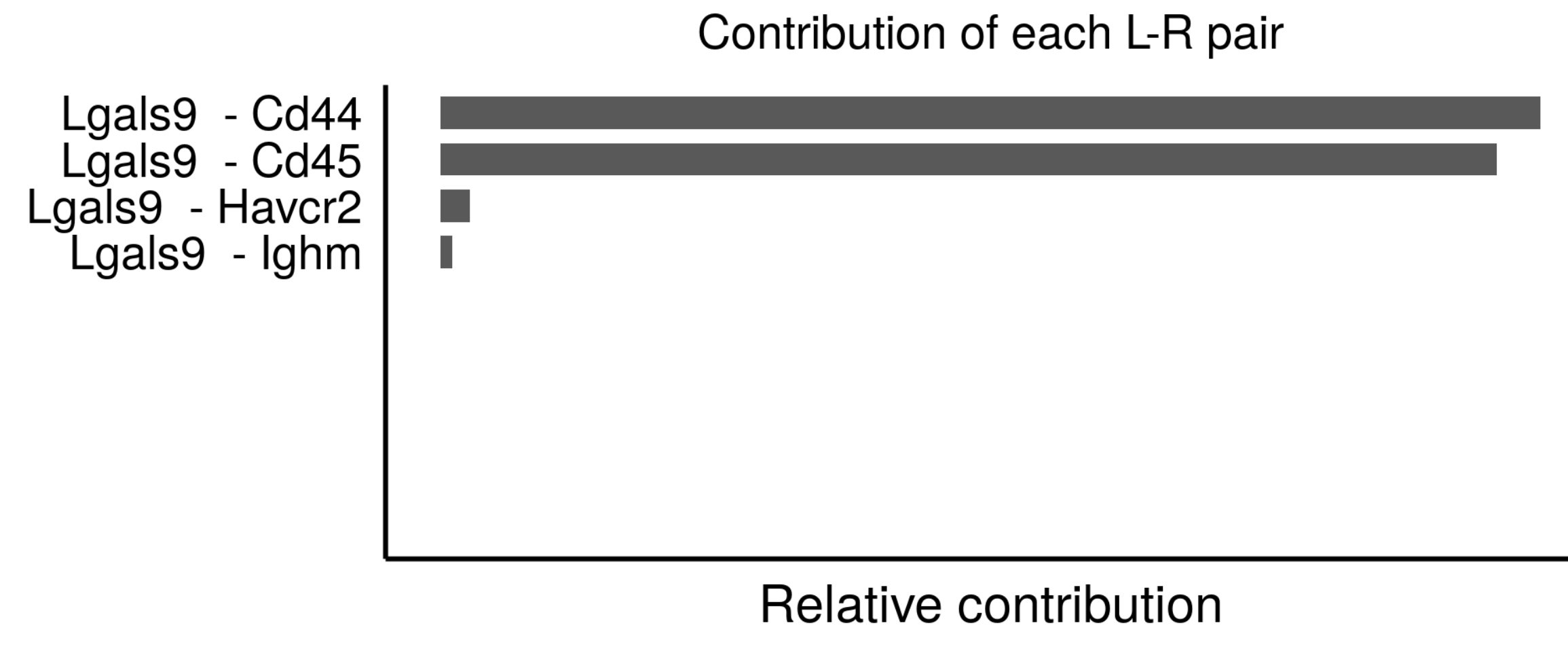

**a**

**b**

**c**

**d**

**e**

**figure S2. ScRNA-seq signaling responses during infection**. (a) KEGG enrichment analysis of upregulated activity for neutrophil subpopulations identified via scRNA-seq at 7 DPI in mouse skin lesion. (b) UMAP plot of *Il17a* and *Il17f* expression by cells in infected skin lesion at 7 DPI vs in uninfected skin. Orange color represents cells with increased gene expression. (c) Cellchat heatmap of CD80 signaling pathway network for cell types from infected skin at 7 DPI. (d) Cellchat heatmap of Galectin signaling pathway network for cell types from infected skin at 7 DPI. (e) Cellchat heatmap of CD80 signaling pathway network for cell types from infected skin at 7 DPI. (f) Contribution of each ligand receptor pair in the GALECTIN signaling network to the overall signaling pathway at 7 DPI.

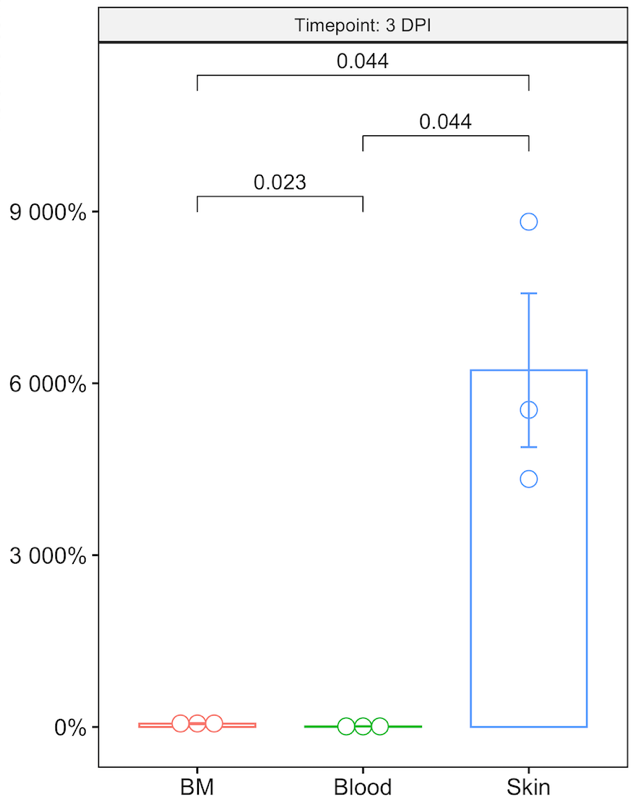

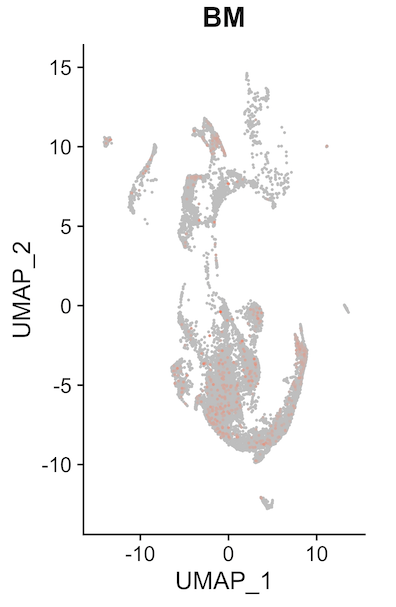

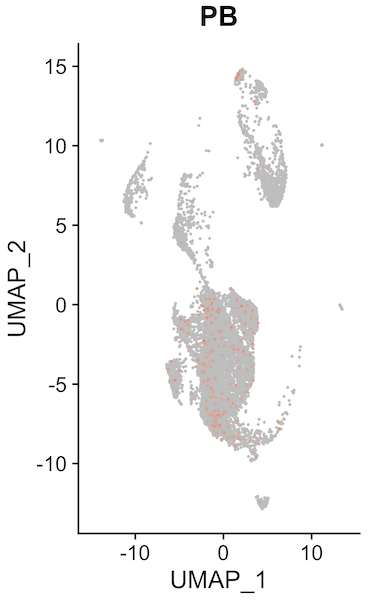

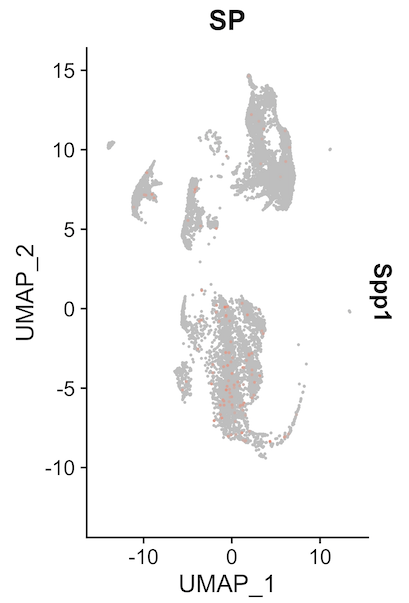

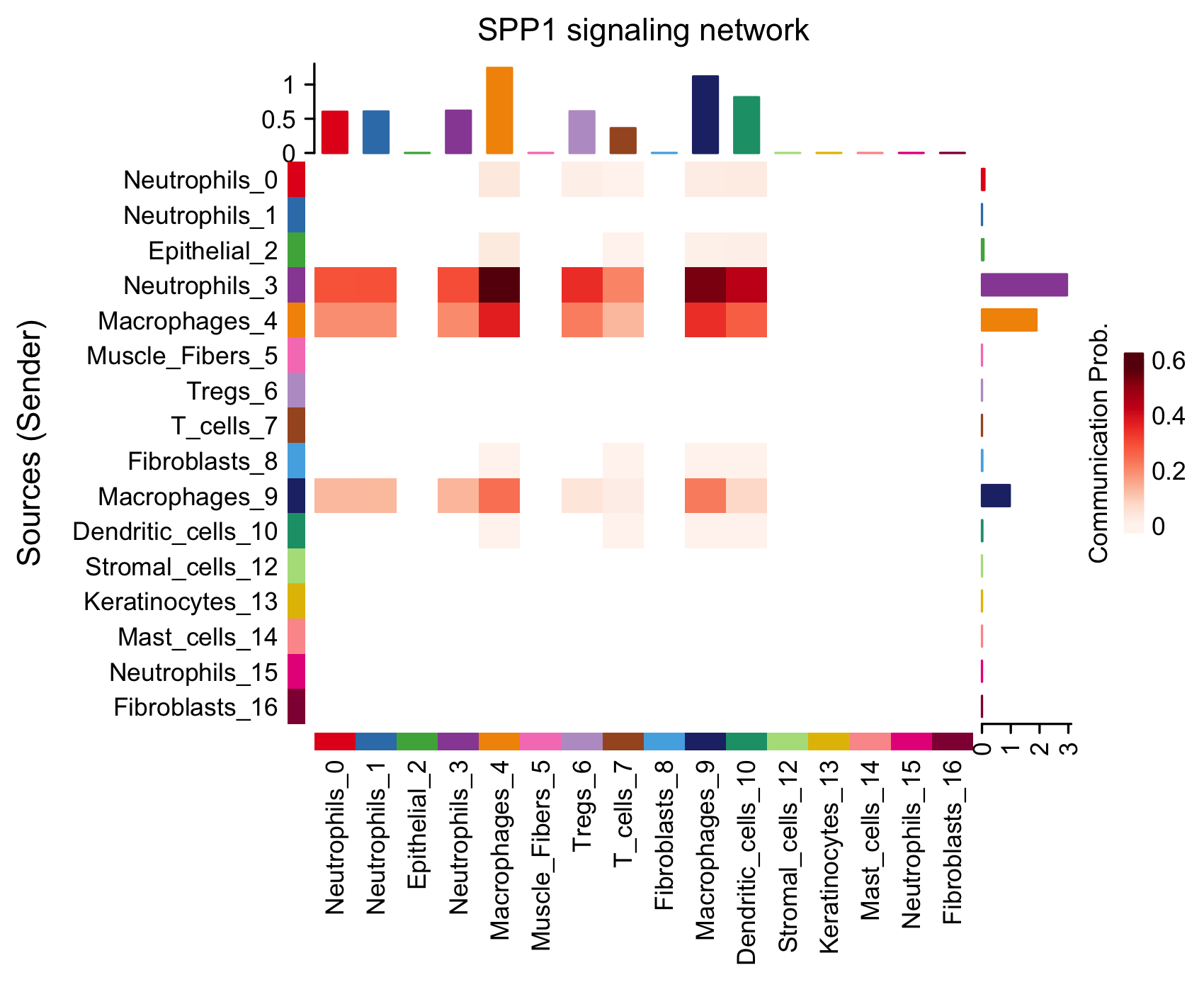

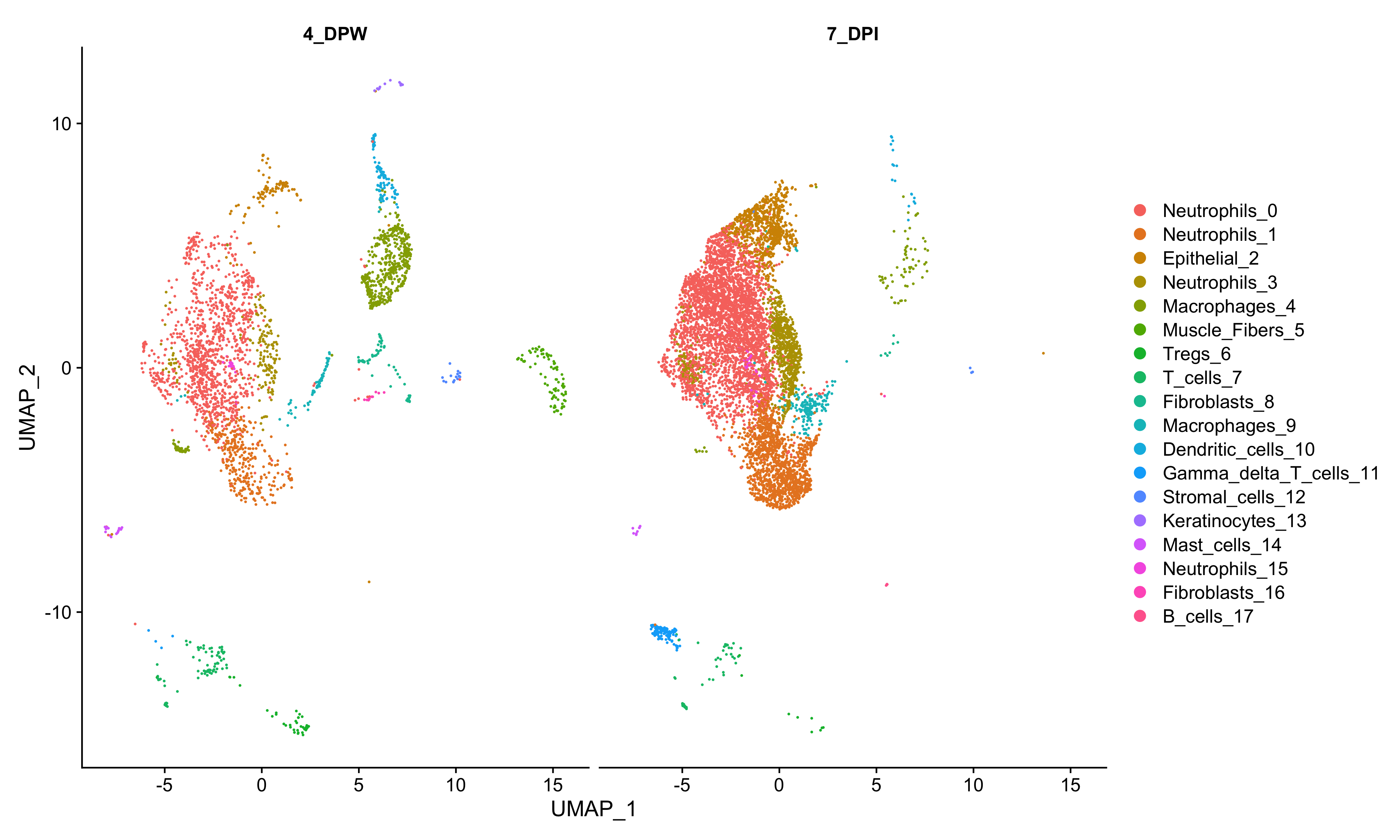

**a**

**b**

**c**

**e**

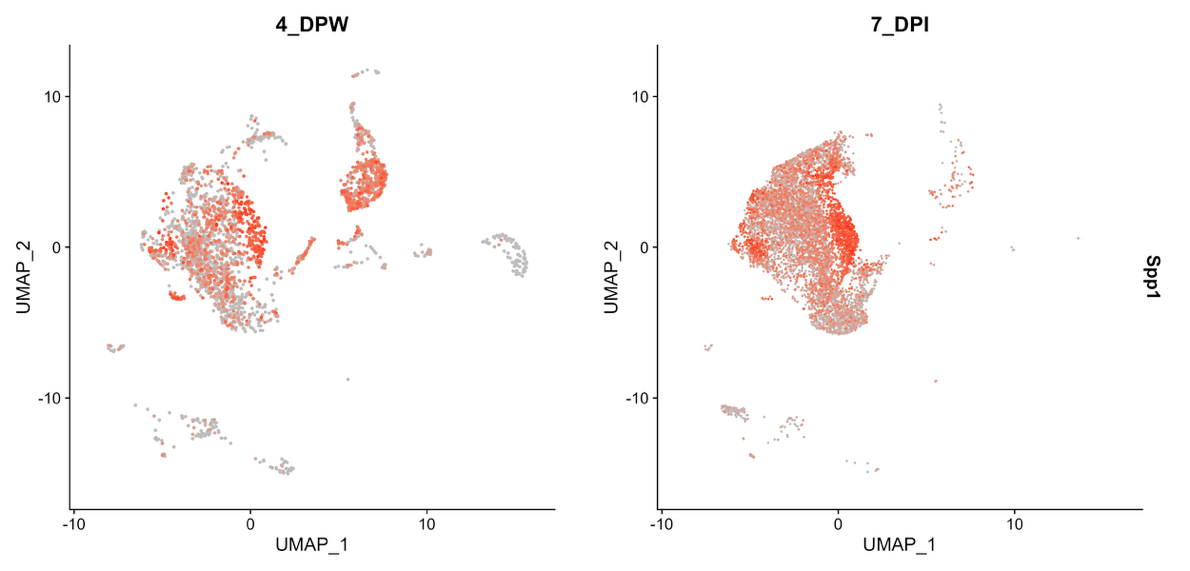

**d**

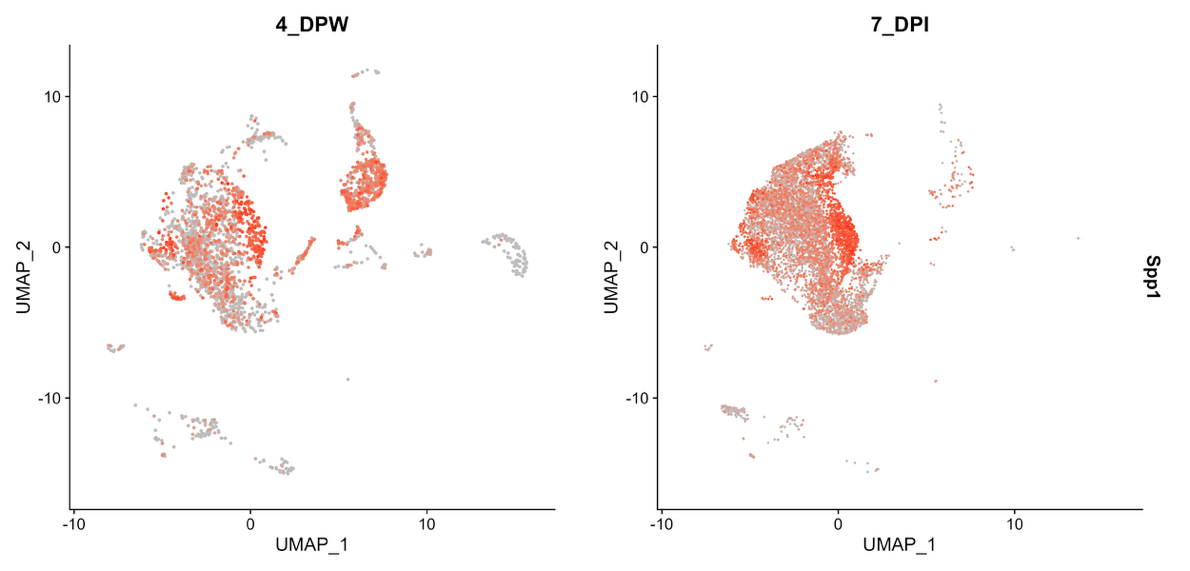

**figure S3. Comparative analysis of Spp1 in different tissues and conditions**. (a) UMAP plots from reanalysis of Xie et al existing dataset showing Spp1 expression levels in neutrophils isolated from bone marrow (BM), peripheral blood (PB), and spleen (SP). (b) Relative mRNA expression of *Spp1* in neutrophils isolated from the BM, blood, and skin following dermonecrotic *S. aureus* skin infection, GAPDH was used as an internal control. (c) UMAP showing cell types from comparative analysis of our 7 DPI scRNA-seq data and existing scRNA-seq dataset of skin at 4 days post wounding (DPW) from Vu et. Al. (d) UMAP plots showing *Spp1* expression levels in infected vs wounded skin from analysis in (c). (e) Cellchat heatmap of *Spp1* signaling senders and receivers present at 4 days post wounding.

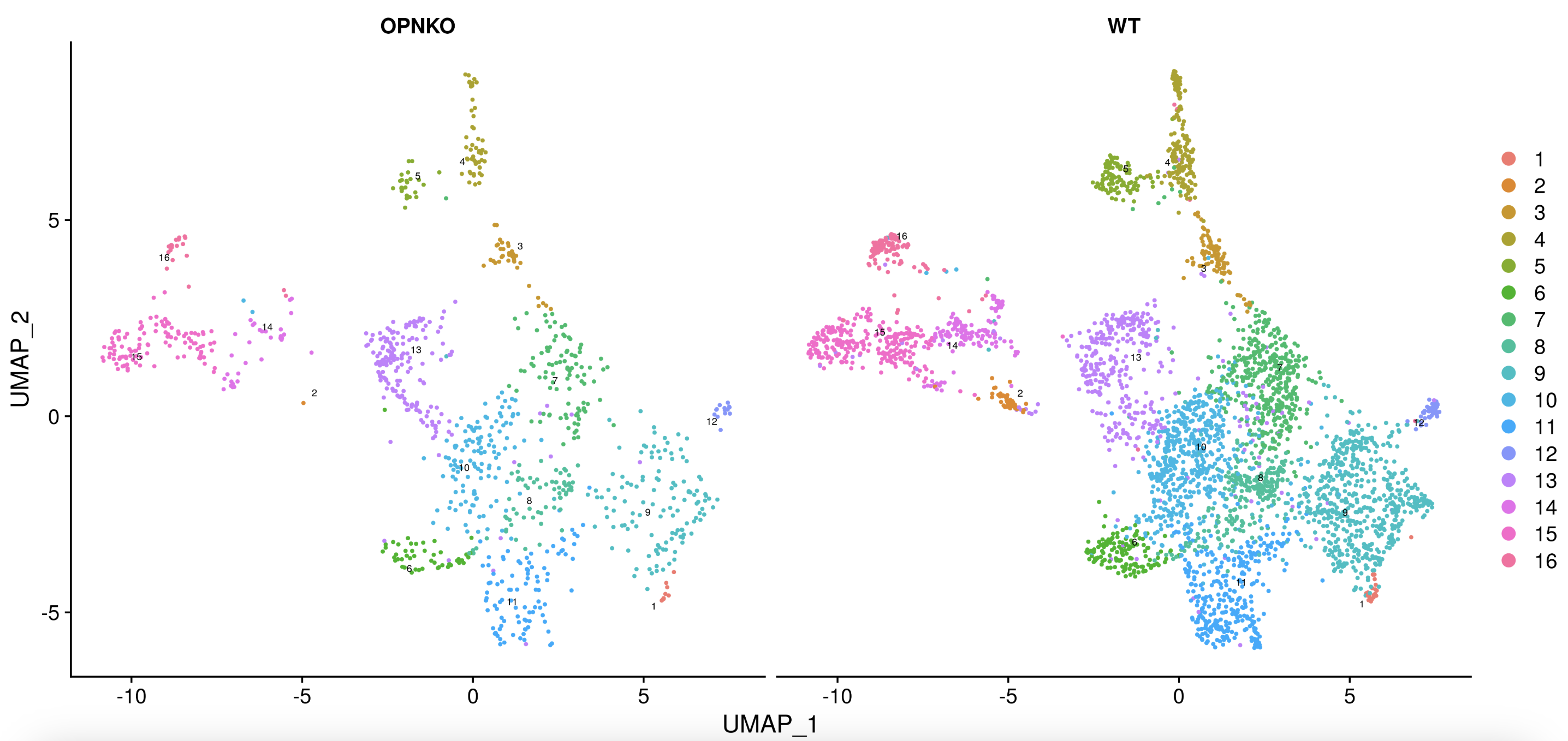

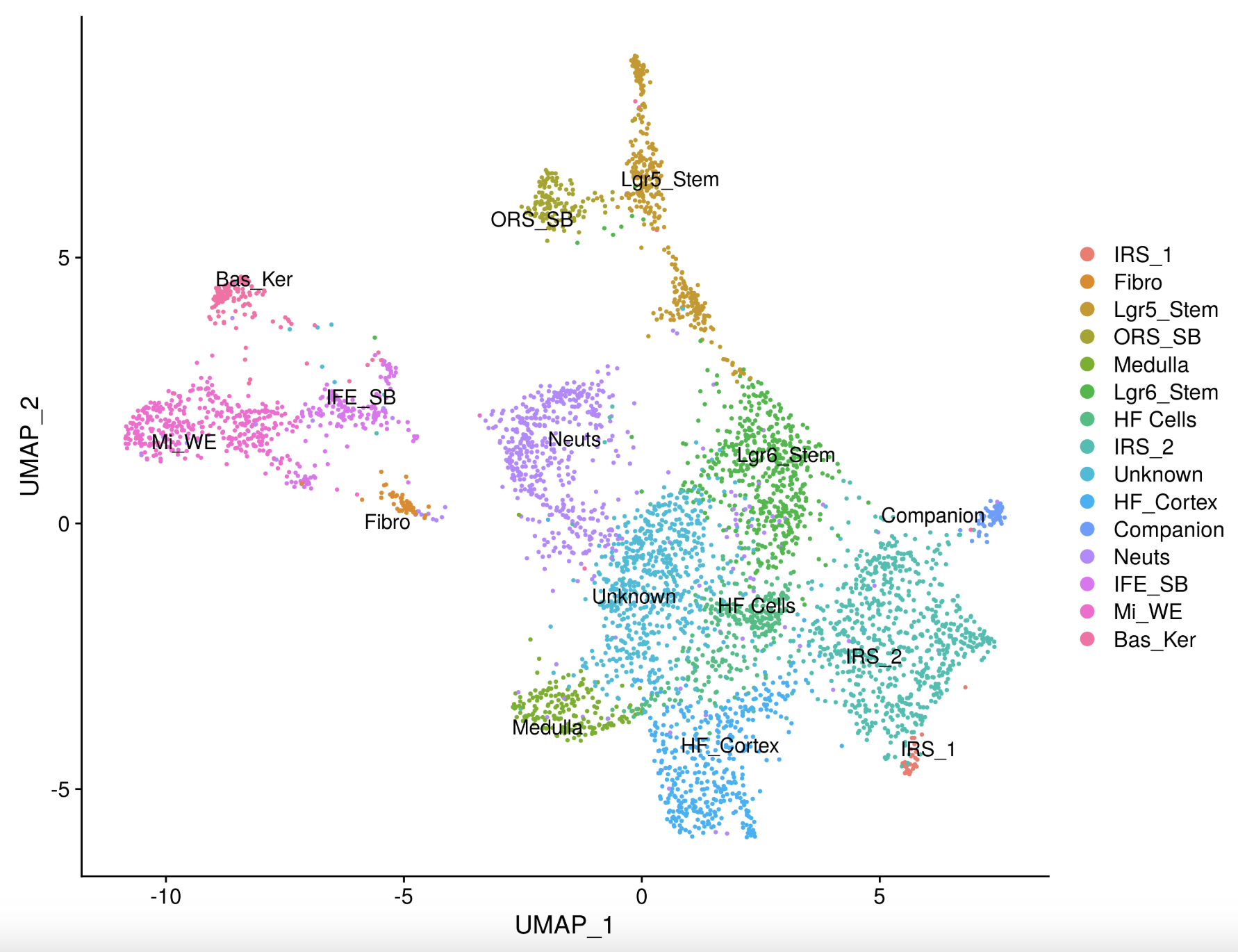

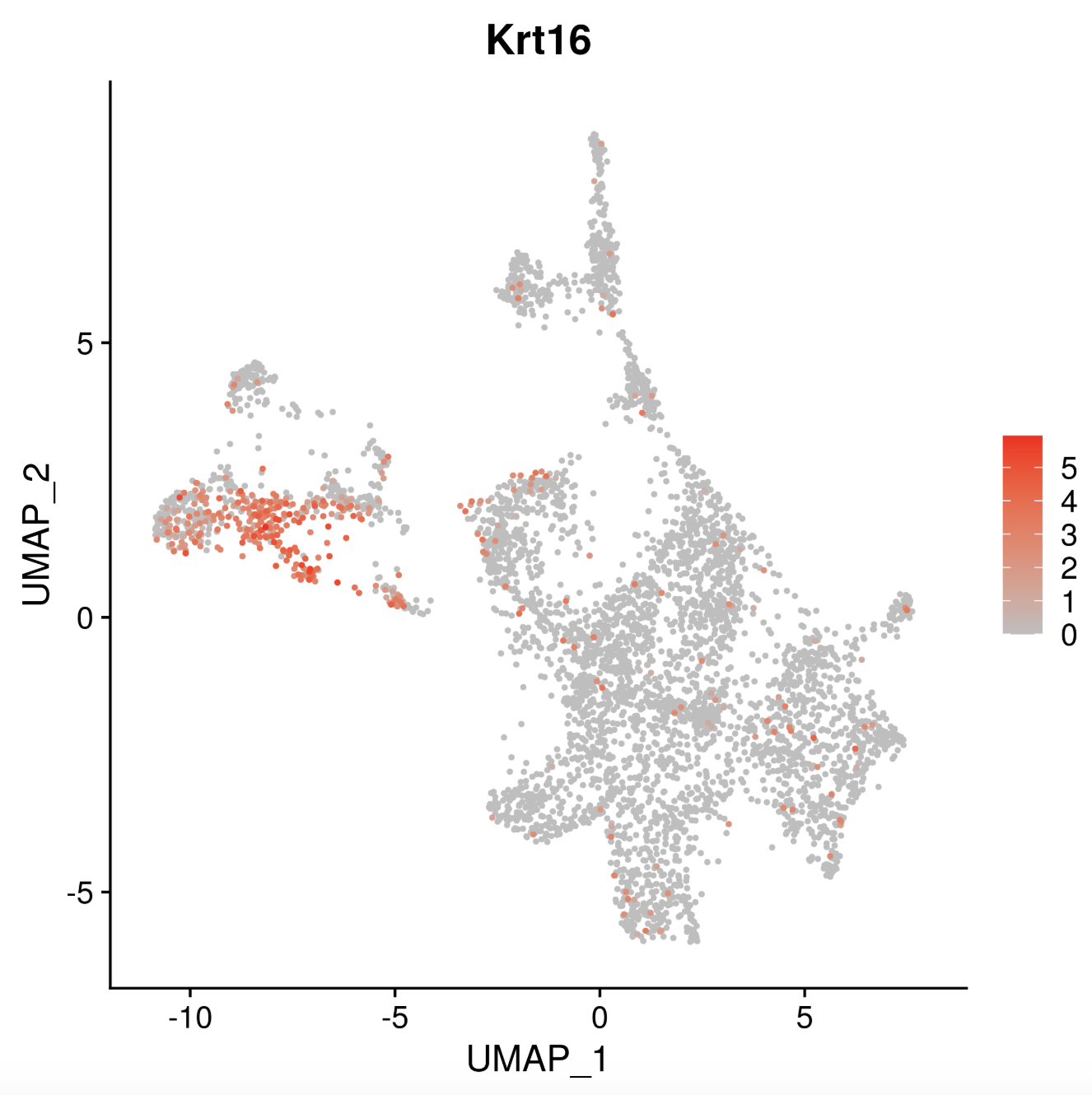

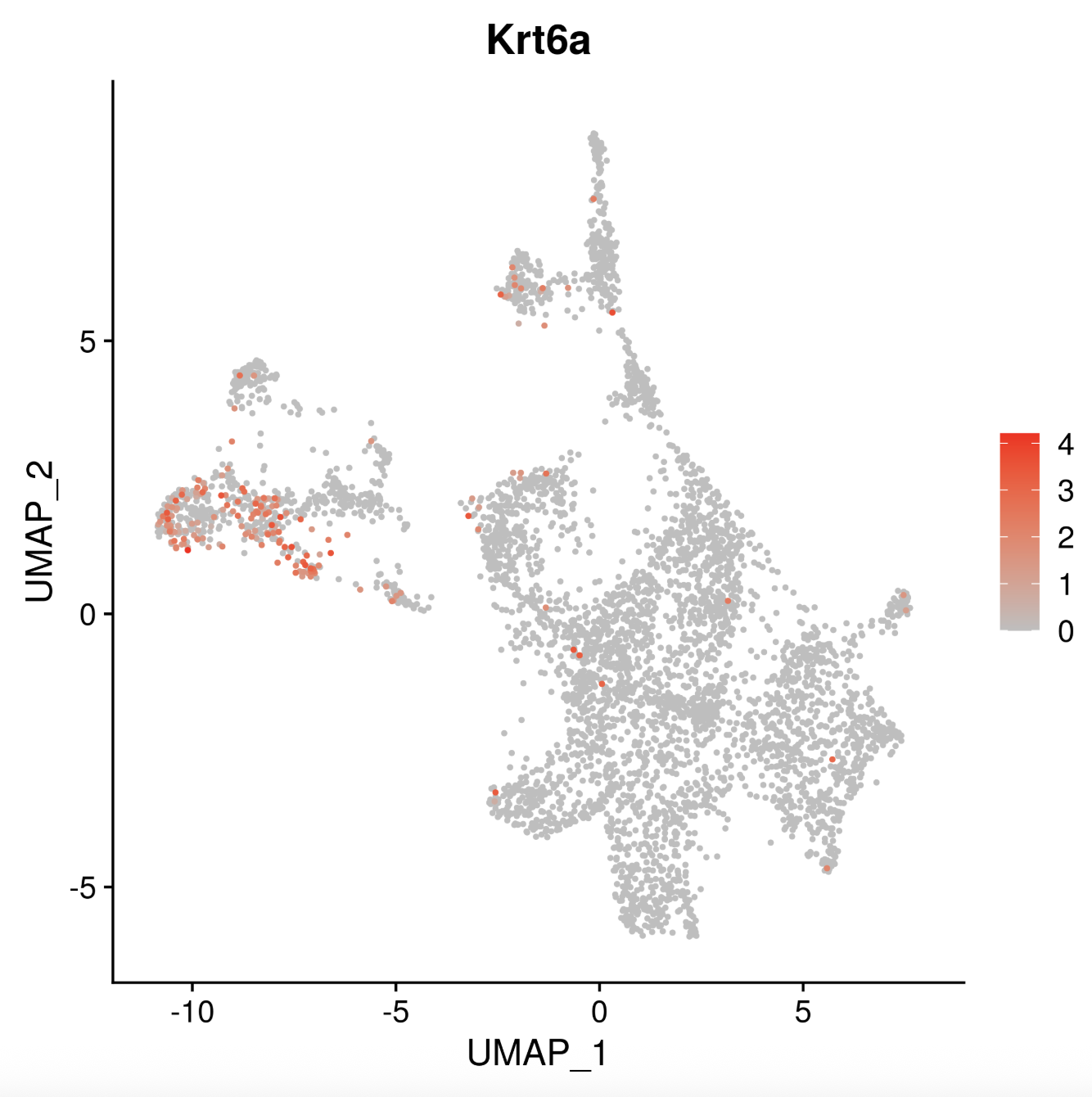

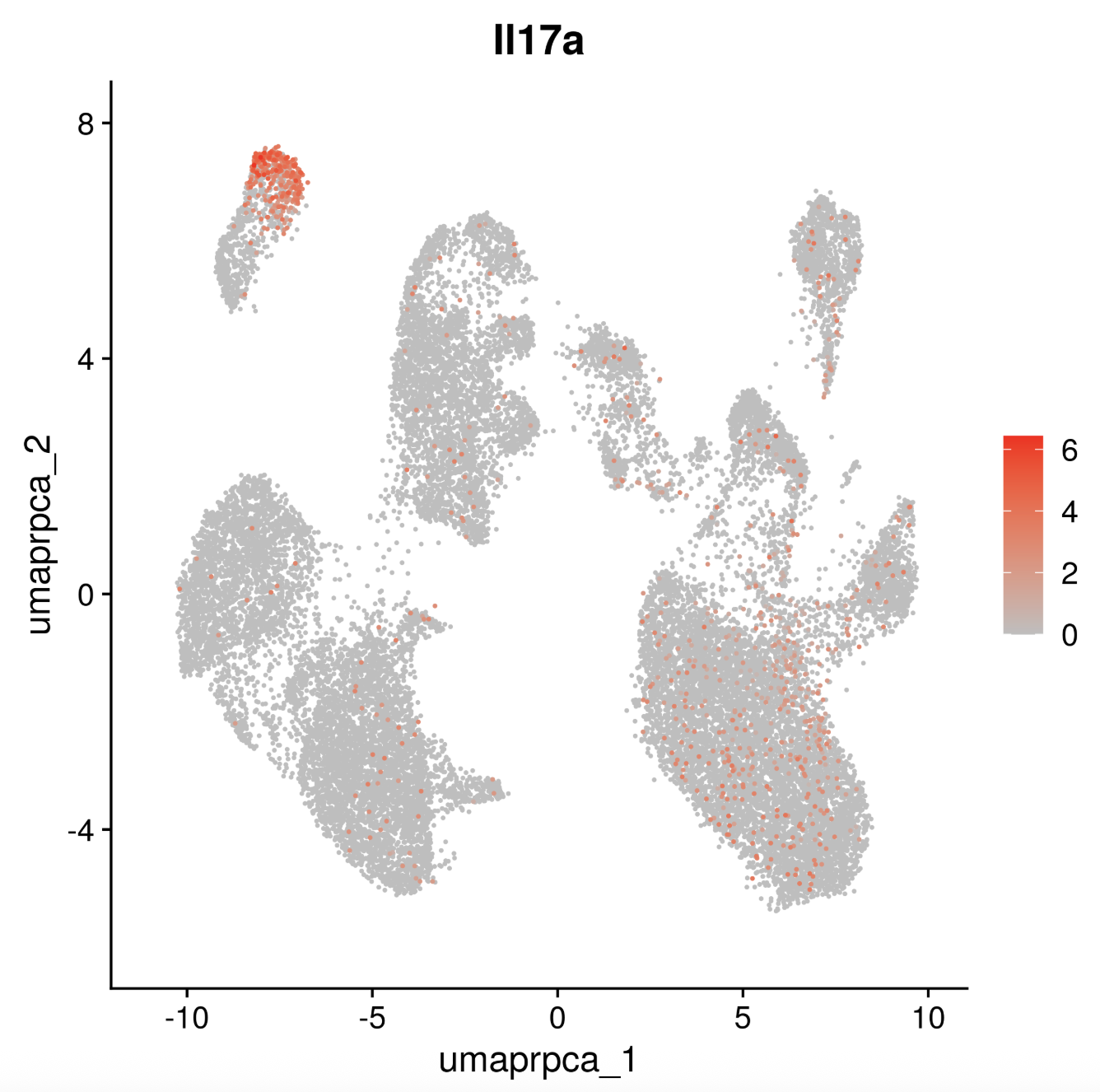

**a**

**b**

**d**

**e**

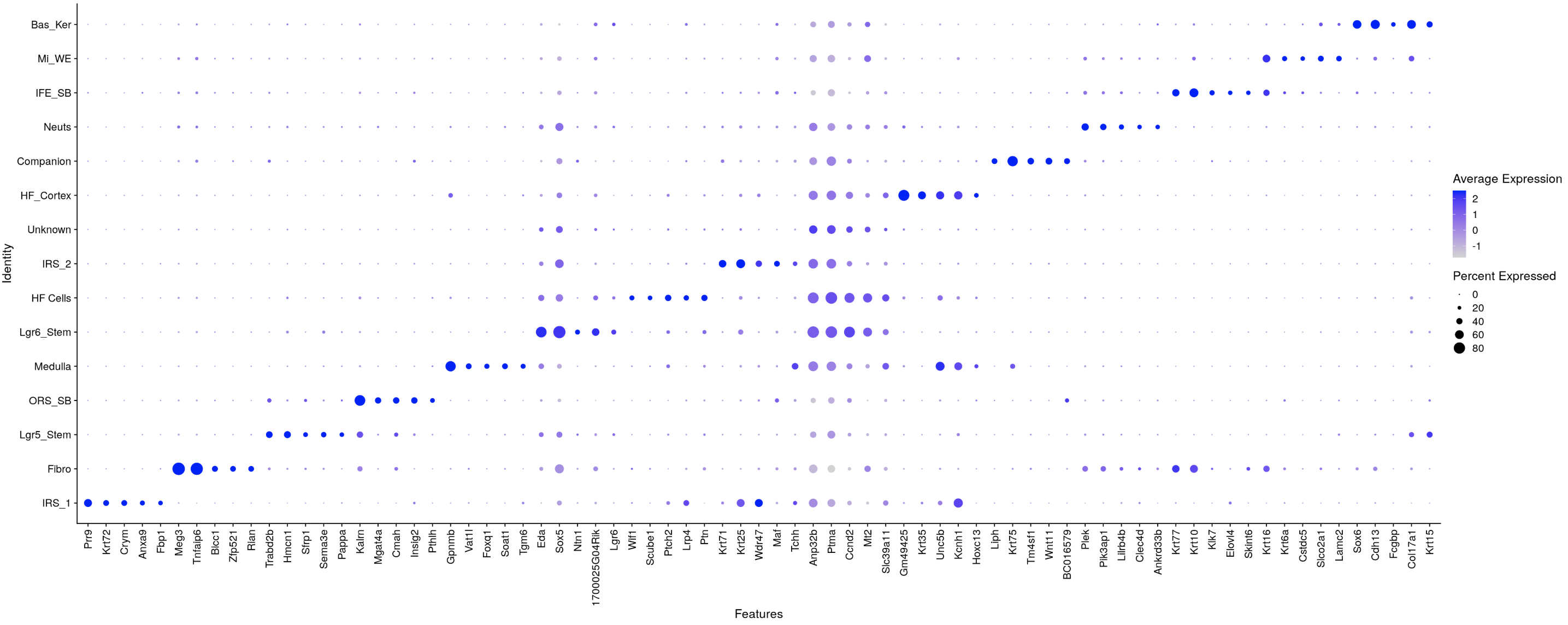

**c**

**figure S4. OPN KO and WT gene expression and keratinocyte subclustering**. (a) UMAP *IL17a* expression levels are high in gd T cells (b) Dotplot of representative genes used to identify cell types following subclustering of keratinocytes from OPN KO and WT mice scRNA-seq data. (c) UMAP of scRNA seq subclustering of keratinocytes shows cluster distribution in OPN KO vs WT mice(d) UMAP plot showing identified cell types from keratinocyte subclustering based on genes in (b). (e) UMAPs of *Krt16* and *Krt6b* expression in keratinocyte subtypes show high levels in migrating wound edge (Mi_WE) population from (d).

**f**
